# Common gene signature model discovery and systematic validation for TB prognosis and response to treatment

**DOI:** 10.1101/2022.11.28.518302

**Authors:** Roger Vargas, Liam Abbott, Nicole Frahm, Wen-Han Yu

**Affiliations:** Bill & Melinda Gates Medical Research Institute, Cambridge, MA, USA; Harvard University, Cambridge, MA, USA

## Abstract

While blood gene signatures have shown promise in tuberculosis (TB) diagnosis and treatment monitoring, most signatures derived from a single cohort may be insufficient to capture TB heterogeneity in populations and individuals. Here we report a new generalized approach combining a network-based meta-analysis with machine-learning modeling to leverage the power of heterogeneity among studies. The transcriptome datasets from 57 studies (37 TB and 20 viral infections) across demographics and TB disease states were used for gene signature discovery and model training and validation. The network-based meta-analysis identified a common 45-gene signature specific to active TB disease across studies. Two optimized random forest regression models, using the full or partial 45-gene signature, were then established to model the continuum from *Mycobacterium tuberculosis* infection to disease and treatment response. In model validation, using pooled multi-cohort datasets to mimic the real-world setting, the model provides robust predictive performance for incipient to active TB risk over a 2.5-year period with an AUROC of 0.85, 74.2% sensitivity, and 78.3% specificity, which approximated the minimum criteria (>75% sensitivity and >75% specificity) within the WHO target product profile for prediction of progression to TB. Moreover, the model strongly discriminates active TB from viral infection (AUROC 0.93, 95% CI 0.91-0.94). For treatment monitoring, the TB scores generated by the model statistically correlate with treatment responses over time and were predictive, even before treatment initiation, of standard treatment clinical outcomes. We demonstrate an end-to-end gene signature model development scheme that considers heterogeneity for TB risk estimation and treatment monitoring.

**AUTHOR SUMMARY:** An early diagnosis for incipient TB is a one of the key approaches to reduce global TB deaths and incidence, particularly in low and middle-income countries. However, in appreciation of TB heterogenicity at the population and individual level due to TB pathogenesis, host genetics, demographics, disease comorbidities and technical variations from sample collecting and gene profiling, the responses of the molecular gene signatures have showed to be associated with these diverse factors In this work, we develop a new computational approach that combines a network-based meta-analysis with machine-learning modeling to address the existing challenge of early incipient TB prediction against TB heterogenicity. With this new approach, we harness the power of TB heterogeneity in diverse populations and individuals during model construction by including massive datasets (57 studies in total) that allow us not only to consider different confounding variables inherited from each cohort while identifying the common gene set and building the predictive model, but also to systematically validate the model by pooling the datasets to mimic the real-world setting. This generalized predicting model provides a robust prediction of long-term TB risk estimation (>30 months to TB disease). In addition, this model also demonstrates the utility in TB treatment monitoring along with Mycobacterium tuberculosis elimination.

## INTRODUCTION

Tuberculosis (TB) is a leading cause of death globally from the infectious disease agent, *Mycobacterium tuberculosis* (*Mtb*), causing an estimated 10 million new cases of disease and 1.3 million deaths per year (*1*). About 85% of individuals who develop active TB disease (ATB) can be successfully treated, but not all cases are diagnosed (*1*). Better diagnostic tools are necessary to reach the WHO 2035 targets of a 95% reduction in global TB deaths and a 90% reduction in global TB incidence (compared to the 2015 baseline), in addition to developing new interventions for TB prevention and treatment (*1*). An accurate, rapid, point-of-care (PoC), non-sputum-based diagnostic test is required in low and middle-income countries to identify individuals with ATB who require treatment, and to predict which individuals with latent tuberculosis infection (LTBI) are likely to progress to ATB, and therefore require preventive therapy (*2–4*). Further, such a test may be used to monitor treatment response in individuals who commence TB drug treatment to ensure that they are cured at the end of treatment, or to allow for treatment modification if cured before the completion of the standard 6-month treatment regimen (*1*).

Several whole blood diagnostic gene signatures based on host immune responses have demonstrated the ability to accurately distinguish between ATB and healthy controls (HC), as well as ATB and disease manifestations such as other pulmonary diseases (OD) or LTBI (*3, 5–16*). Further, some studies have also demonstrated the use of gene signatures in predicting progression from LTBI to ATB (*3, 5, 11, 13, 16–29*) and in monitoring treatment response (*3, 8, 16, 30–35*). For TB gene signature discovery, most studies utilized a conventional approach beginning with a single cohort with differential gene expression analysis between ATB and different clinical conditions (LTBI, OD or HC), followed by multivariate modeling to define the gene signature. The limited sample size in a single cohort, which is substantially smaller than the number of measured genes, increases the risk of multicollinearity and model overfitting. As a result, the conventional approach often insufficiently captures cohort heterogeneity and therefore generates a gene signature specific to the cohort examined. A systematic comparison of 16 published gene signatures, most of which compared cohorts derived from one location, demonstrated a limited overlap of the genes among these signatures (*2*). Whether this stems from limited heterogeneity in single cohort samples or due to the diversity in the analytical approaches remains unclear.

In addition to diverse immune responses associated with different states in a wide spectrum of *Mtb* infectious pathophysiology (*36*), heterogeneity in cohort studies due to demographics, disease comorbidities, sample collection (whole blood vs. PBMCs) and transcriptomic profiling technologies (qRT-PCR, microarray, or RNAseq) make it challenging to translate the use of any one gene signature into a generalizable PoC diagnostic test for use in real-world patient populations (*2, 18, 37*). Among all existing gene signatures, two models (Sweeney 3 and RISK 6) demonstrated their utility as a screening or triage test for TB diagnosis in multicenter prospective studies (*32, 38*), which showed potential to be more generalizable for different patient populations. To integrate population heterogeneity as part of the model building process, we conducted a meta-analysis where the transcriptome datasets from 37 published studies, all of which contained ATB and at least one other disease conditions (LTBI, OD, or HC), were pooled together to capture various confounding factors inherited from individual studies in order to mimic a real-world setting. The distributions of TB scores in Sweeney 3 and RISK 6, when applied to this pooled dataset, were shown to be less distinguishable between ATB and other disease conditions than when applied in their original populations (**Supplementary Fig. 11E and G**). This poses a challenge for calling a positive case from a screening or triage test for disease diagnosis due to the impact of patient and population level heterogeneity.

To this end, we aimed to assess whether we could leverage the power of individual/population heterogeneity from a collection of 27 transcriptome datasets to establish a generalized multivariate model for TB risk estimation and treatment monitoring. In the process of model development, we first developed a network-based meta-analysis that utilized the collected cohorts containing different clinical/genetic/technical covariates to identify a set of common genes that robustly distinguish ATB from LTBI, HC, OD and ATB treatment across the cohorts. Importantly, the network-based approach was designed to capture not only the most differentiated genes, but also the covariation between genes that elucidated functionally important biological processes that underlie progression to ATB. This approach led to the identification of a biologically relevant and statistically robust gene signature. We then trained two optimized machine-learning (ML) regression models (with full and reduced gene sets) to connect the expression patterns of the gene signature and the dynamic continuum of *Mtb* infection to ATB disease. A systematic validation of the model is presented using 10 independent longitudinal studies to evaluate the potential utility of our model in a real-world setting and 20 additional viral infection studies for testing model specificity to TB. A probabilistic model to estimate TB disease risk is also established for use as a TB screening tool.

## RESULTS

### Development of a network-based meta-analysis approach to identify a common gene signature specific to active tuberculosis

First, we aimed to identify a set of common genes consistently differentially expressed in ATB cases relative to other conditions, including HC, LTBI, OD and undergoing TB treatment (Tx), while accounting for cohort/population heterogeneity among the studies. To do so, we developed a network-based meta-analysis approach where a gene covariation network was established to consider genes not only differentially expressed but also co-varied among the studies based on the meta-analysis (**Fig. 1A**). We identified a TB-specific common gene set and used it as the training dataset to build the ML predictive model for TB risk estimation and treatment monitoring. Specifically, we collected whole blood transcriptome data from 27 published studies (the discovery dataset) in which each dataset had ATB and at least one other contrasting condition for a given cohort of individuals (**Supplementary Table 1**). For each contrasting condition comparison (e.g., ATB vs. HC) we conducted a differential gene expression analysis within a cohort resulting in a list of differentially expressed genes with corresponding log fold-changes (logFC) (**Fig. 1A1**). After repeating the analyses across cohorts, we created an ***N*** × ***M*** matrix ***X*** with logFC values for ***M*** cohorts and ***N*** genes (**Fig. 1A2**). We further simplified ***X*** by converting the matrix values to +1, 0, −1 to account for uncertainty of the data, resulting in a matrix that captured differentially expressed genes that were either upregulated or downregulated between ATB and the other conditions (HC, LTBI, OD or Tx). To capture co-varying genes across cohorts, we constructed a gene covariation network from the matrix ***X*** where each node represented a gene, and the edge weight between nodes was calculated by dot product between the paired genes in the matrix ***X***. The network only retained edges with weights ≥ 3, which was determined to represent significant covariation from permutation tests (**Fig. 1A2, Supplementary Fig. 1**). The weighted degree of each node as the sum of the weights of the edges assigned to the node was also calculated to infer its node centrality (**Supplementary Fig. 2**). The closer to the center (the higher weighted degrees) the nodes are, the more consistently the genes respond to active TB among various studies. To define a robust gene signature discriminating ATB from different conditions, we built four covariation networks for each condition comparison and rank ordered genes based on their centrality (**Fig. 1A3**). Finally, we identified a set of 45 genes that were in the top 5% of central genes from the weighted degree distribution of the network, were central in at least two networks, and had average logFC ≥ 0.5 or ≤ −0.5 across the studies (**Supplementary Fig. 3**).

**Figure 1.**
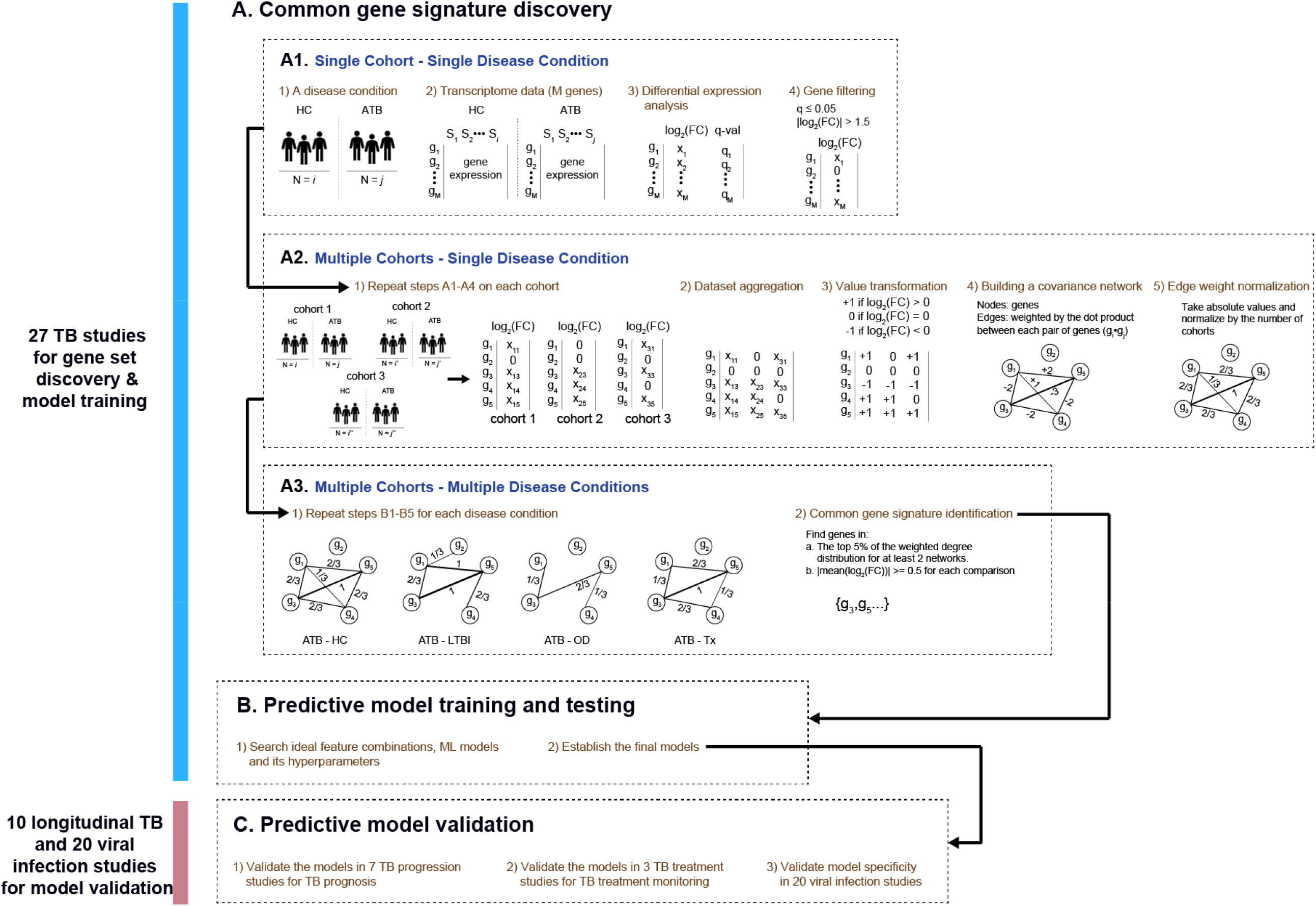
An end-to-end gene signature model development scheme. (A) The network-based meta-analysis approach. (A1) Schematic of differential gene expression calculations for a single cohort given a disease condition. (A2) Steps for integrating differential gene expression results for multiple cohorts into a gene covariation network. (A3) Four different networks were constructed corresponding to four different disease conditions: ATB v HC, ATB v LTBI, ATB v OD, ATB v Tx. From these four networks, the top weighted nodes across networks were selected to form the common ATB-specific gene signature. (B) ML predictive model training and testing based on the common gene signature. (C) Steps for the predictive model validation.

We validated this gene set with a volcano plot showing the average logFC across datasets against the weighted degree of each gene for each network constructed and highlighted the 45 gene set (**Fig. 2A-D**). Consistent with our selection strategy, we find that most genes in this gene set are highly differentiated when averaged across cohorts for each disease comparison and heavily weighted (centered) in most of the networks. In addition, we plotted differential expression of the 45 genes in heatmaps across all datasets (**Fig. 2E-H**) and observed that gene patterns of upregulation and downregulation were consistent across cohorts within each disease condition and between disease conditions. Genes upregulated in ATB vs. HC for example were also upregulated in ATB vs. OD, demonstrating a consistent signal for this gene set across cohorts within and between disease conditions. However, a few genes (such as HP, AEACAM1) showed slightly inconsistent expression patterns in ATB vs. OD.

**Figure 2.**
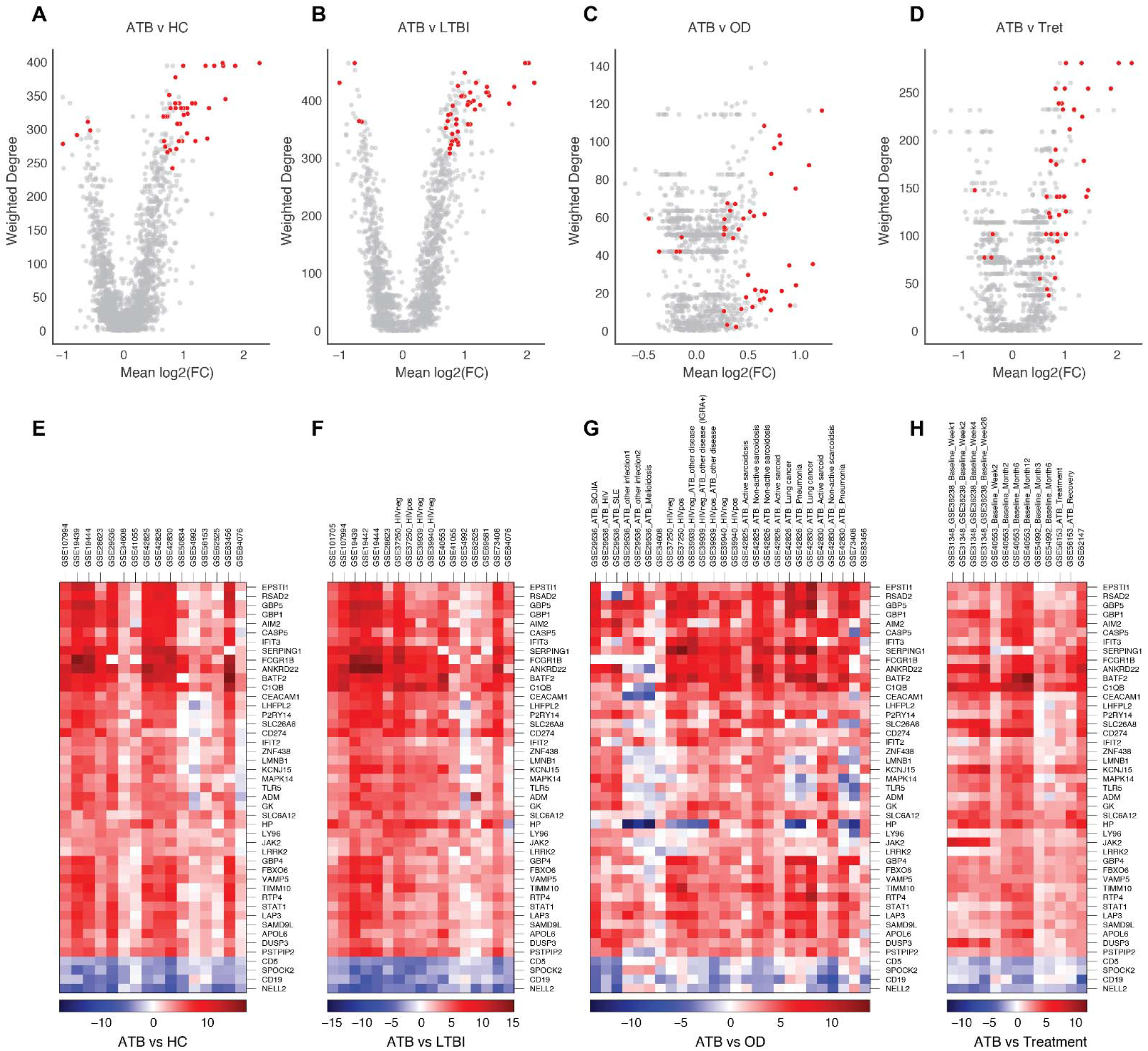
Network analysis results and gene signature. (A-D) Volcano plots displaying the mean log2(Fold Change) of gene expression across cohorts against the weighted degree for each network for all genes within each network. Forty-five candidate genes highlighted in orange within each plot. (E-H) Heatmaps of log2(Fold Change) for each of the 45 candidate genes (rows) between disease conditions displayed for each cohort (columns) that were used in constructing each network. The corresponding GSE ID of each cohort was labeled on columns. If a cohort contains multiple clinically defined populations, a specific comparison is also highlighted on the column.

We next sought to identify a biological interpretation for the 45-gene set. First, we performed a gene set enrichment analysis and found an enrichment of genes involved in immune function such as interferon gamma (IFN-*γ*) and alpha/beta (*α/β)* signaling, IL-6 signaling and Toll-like receptor cascades (**Supplementary Table 2**). Next, we mapped all 45 genes in the protein-protein association network using the STRING database (*39*). Strikingly, we found 71% (32/45) of the genes were interconnected (**Supplementary Fig. 4**). The connections between genes may indicate physical interaction between some of the encoded proteins, or that the proteins jointly contribute to a shared function (*39*). Proteins IFIT2 and IFIT3 centered in the network are IFN-α/β induced proteins and highly expressed during cell-intrinsic immune response to *Mtb* infection (*40*). Further, we observed that STAT1 forms associations with many proteins in the network (*n* = 17); STAT1 is known to be an immunoregulatory factor that regulates both IFN-α/β and IFN-γ-mediated responses (*7, 41, 42*). Taken together, this report using a network-based meta-analysis approach on whole blood transcriptome datasets provided the first systematic evidence to identify a set of 45 genes that commonly appeared in all 27 studies and enriched for genes actively involved in interferon activation responding to infection.

### Machine-learning (ML) model building and optimization

The scope of the ML work was to generate a TB score, a continuous variable, with a fixed range between 0 and 1, representing a dynamic continuum from *Mtb* infection to disease as well as following disease treatment back to cured. Multivariate regression was then used to establish the relationship between outcome (TB disease states) and the expression patterns of the ATB-specific gene signature identified from our network-based meta-analysis (**Fig. 1B)**. To capture heterogenous responses of individuals in different TB states among the cohorts, the whole discovery dataset (27 studies) was pooled together (**Supplementary Table 1**), and the ML model regressed to two states: 1 (ATB) and 0 (HC, LTBI, Tx or OD). The ML framework began with feature selection where the features fed into ML models were derived from an expression ratio between any pair of the common genes. Feature selection underwent two steps: univariate and multivariate feature selection. Combination of two steps allowed removal of irrelevant or redundant features from the original 990 gene-paired features down to 41 features (**Supplementary Table 4**) and mitigated the risk of model overfitting while improving predictive performance. Next, we assessed the best models among 7 ML regression algorithms using the discovery dataset based on 5-repeated 5-fold nested CV framework. Random forest (RF) model consistently outperformed other 6 ML models based on the performance metrics (average outer cross-validation R^2^ = 0.51, MSE = 0.11, AUROC = 0.91), while multilayer perceptron model, an artificial neural network approach, was second-best (average outer cross-validation R^2^ = 0.48, MSE = 0.11, AUROC = 0.90) (**Supplementary Table 3, Supplementary Fig. 6**). We therefore generated a final RF model trained using the full discovery dataset; its parameters are listed in **Supplementary Table 4**. Given that blood transcriptomic signatures have held great promise for development of screening and triage tests (*30, 32, 38*), we further adapted a stability selection approach to perform a sensitivity analysis and downselected the most robust features while maintaining prediction performance (**Supplementary Fig. 7**). Using a forward stepwise selection within a 5-fold nested CV framework, we evaluated the RF algorithm by adding one feature at a time starting from the top ranked features. The final model with 12 features (paired genes) was determined with slightly reduced performance (average outer cross-validation AUROC = 0.85) (**Supplementary Fig. 8**). As a result, two models, labeled as the full and reduced model, were generated, and listed in **Supplementary Table 4**. Importantly, among the 18 genes selected by the reduced model, 8 out of 18 genes (ANKRD22, APOL6, BATF2, DUSP3, FCGR1B, GBP4, GBP5, and VAMP5) were repeatedly identified by the best performing gene signature models for incipient TB diagnosis reported previously (*43*) (**Supplementary Fig. 5**), in which DUSP3 and GBP5 included in Sweeney 3 gene signature have been tested in a PoC setting for TB diagnosis (*5, 38, 44*). Seven additional genes (ADM, CD274, FBOX6, LMNB1, IFIT2, NELL2, and ZNF438) have been identified as part of several gene signatures associated with active TB (*7, 28, 45, 46*). For the remaining 3 new genes (CD5, LRRK2 and SPOCK2), CD5-expressing regulatory B cells were found to negatively regulate Th17 and to associate with active TB (*47*). SPOCK2 expression was shown to be associated with lung injury induced by viral infection (*48*) and the mouse LRRK2 knockout model suggested that LRRK2 regulates innate immune responses and lung inflammation during *Mtb* infection (*49*). Taken together, we demonstrated that our gene signature discovery approach is robust enough to not only produce comparable results to other models but to expand on previous models with better performance.

### Validation of the gene signature models in TB disease risk prediction

To systematically validate the models, we collected 10 independent longitudinal studies (7 TB progression and 3 TB treatment cohorts) from multiple countries and different target populations (**Fig. 1C**) (**Supplementary Table 1**). We first pooled all 7 TB progression datasets (*17–19, 27, 28, 31*) and stratified the data using different time intervals to disease and LTBI without progression. TB scores generated from the full model showed a graded increase along time intervals beginning >2 years to disease, and the score distributions from all progression groups were distinguished from the no progression group (**Fig. 3A**). Similar observations were found in the reduced model (**Supplementary Fig. 9**). Furthermore, one of the progression studies (the Leicester household contact study (*31*)) categorized subjects into incipient, subclinical, or clinical TB, according to their clinical phenotypes at the time of sampling. The full model was able to statistically separate the subjects’ scores between clinically defined subgroups and healthy controls as well as in-between clinically defined subgroups (**Fig. 3B**), suggesting that the TB scores derived by the RF model recapitulated host responses to clinical TB pathogenesis. With the same pooled dataset, we compared TB scores generated by the four previously published models, which showed good performance in short-term TB risk estimation (*3, 5, 18, 27*), and similar trends between the scores and time intervals to disease were observed among the models, except Sweeney3, in which the model might be sensitive to different cohort populations, resulting in bimodal distributions found in several groups (**Supplementary Fig. 10**). For most of the published models the score distributions in the groups distant to disease progression (> 12 months) were less discriminating to the no progression group, which imposes a challenge on incipient TB diagnosis at early timepoints (**Supplementary Fig. 10**).

**Figure 3.**
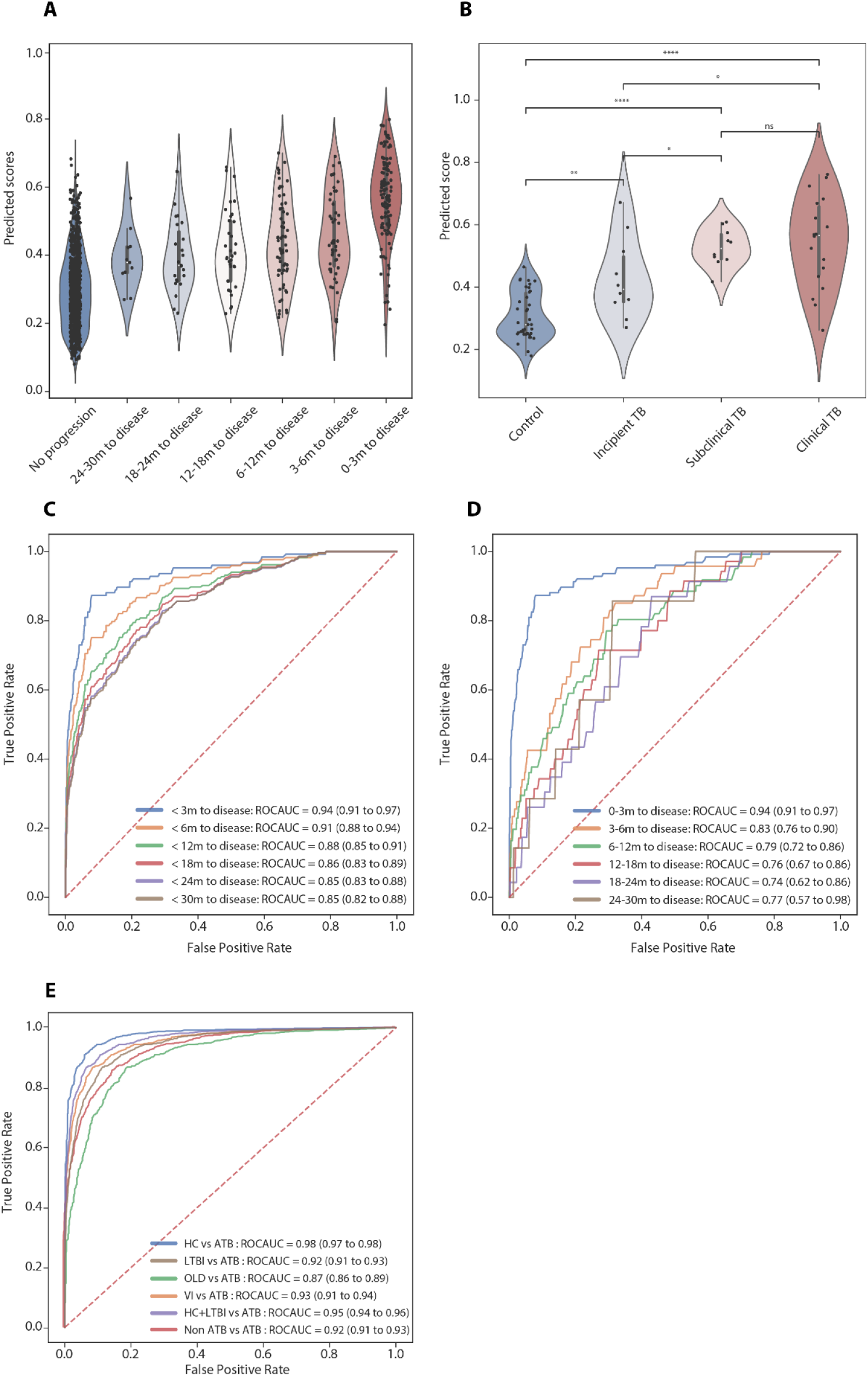
Systematic model validation using TB progression cohorts. The distributions of TB scores generated by the full model, stratified by categorical interval to disease (datapoints *n* = 1281) (A), and clinically defined TB states (datapoints *n* = 137) (B) are shown in the violin plot. Individual datapoints are plotted as a point in the violin plot. P-values were calculated using the Mann-Whitney U test with Bonferroni correction (ns, *p* < 1; *, *p* < 0.05; **, *p* < 0.01; ***, *p* < 0.001; ****, *p* < 0.0001). (C and D) Receiver operating characteristic curves depict diagnostic performance of the model for incipient TB, stratified by time intervals to disease (< 3, < 6, <12, <18, < 24, < 30 months) (C) and mutually exclusive time intervals to disease (0-3, 3-6, 6-12, 12-18, 18-24, 24-30 months) (D). Area under the curve and 95% confidence intervals for each interval to disease are also shown. Diagnostic performance of the model in differentiating between ATB versus HC, LTBI, OLD, and/or viral infection (VI) using a pooled dataset of all 57 collected cohort studies (**Supplementary Table. 1**) (datapoints *n* = 6290) are showed in ROC curves (E).

We next examined prognostic performance of the models by computing AUROCs, stratified by multiple time intervals to disease (< 3, < 6, <12, <18, < 24, < 30 months). Alternatively, mutually exclusive time intervals to disease (0-3, 3-6, 6-12, 12-18, 18-24, 24-30 months) were also examined as a sensitivity analysis. For the full model, AUROC for identification of incipient TB within 3 months was able to achieve 0.94 (95% CI 0.91-0.97) and declined along with increasing interval to disease. Despite that, the AUROC for the group over a 2-year period (< 30 months) was still 0.85 (95% CI 0.82-0.88) (**Fig. 3C, Table. 1**). Interestingly, similar AUROCs were observed in the reduced model even though the model incorporates fewer features (**Supplementary Fig. 9, Table. 1**). Both models outperformed the previously published models in discriminating between progressors and non-progressors in all time periods, except that RISK6 provided a comparable but moderately reduced AUROC result (**Supplementary Fig. 10, Table. 1**). Furthermore, we investigated prognostic performance in each mutually exclusive time period across the models. AUROC values for the full model ranged from 0.94 (0-3 months, 95% CI 0.91 to 0.97) to 0.74 (18-24 months, 95% CI 0.62 to 0.86); values for the reduced model ranged from 0.94 (0-3 months, 95% CI 0.91 to 0.97) to 0.73 (12-18 months, 95% CI 0.63 to 0.82). There was no statistical difference between the two models (**Table 2**). As compared to the previously published models, both of our models had better discrimination between progressors and non-progressors in all mutually exclusive time periods except RISK6, which performed slightly better than our full model 6-12 months prior to disease (**Table. 2**). Remarkably, our models showed great improvement in prognostic performance particularly for the groups distant to disease (12-18, 18-24, and 24-30 months) with AUROCs ranging from 0.74 to 0.77 for the full model and 0.73 - 0.89 for the reduced model compared to range 0.67 - 0.70 for RISK6 (**Table 2**).

**Table 1.** Prognostic performance (AUROC with 95% confidence intervals) of the models developed in this report and published previously on the combined validation datasets for identification of incipient TB within a 2.5-year period, stratified by inclusive time interval to disease.

| Models | < 3m to disease | < 6m to disease | < 12m to disease | < 18m to disease | < 24m to disease | < 30m to disease |
| --- | --- | --- | --- | --- | --- | --- |
| Full model | 0.94 (0.91 to 0.97) | 0.91 (0.88 to 0.94) | 0.88 (0.85 to 0.91) | 0.86 (0.83 to 0.89) | 0.85 (0.83 to 0.88) | 0.85 (0.82 to 0.88) |
| Reduced model | 0.94 (0.91 to 0.97) | 0.89 (0.85 to 0.92) | 0.87 (0.84 to 0.90) | 0.85 (0.82 to 0.88) | 0.85 (0.82 to 0.88) | 0.85 (0.82 to 0.88) |
| Sweeney 3 | 0.68 (0.63 to 0.74) | 0.62 (0.57 to 0.66) | 0.62 (0.58 to 0.66) | 0.60 (0.56 to 0.64) | 0.59 (0.55 to 0.63) | 0.59 (0.55 to 0.63) |
| RISK 6 | 0.92 (0.89 to 0.95) | 0.87 (0.84 to 0.91) | 0.85 (0.82 to 0.89) | 0.83 (0.80 to 0.86) | 0.82 (0.79 to 0.85) | 0.82 (0.79 to 0.85) |
| BATF2 | 0.68 (0.62 to 0.73) | 0.65 (0.60 to 0.69) | 0.62 (0.58 to 0.66) | 0.60 (0.56 to 0.64) | 0.59 (0.55 to 0.63) | 0.58 (0.55 to 0.62) |
| Suliman 4 | 0.90 (0.86 to 0.93) | 0.84 (0.81 to 0.88) | 0.80 (0.77 to 0.84) | 0.77 (0.74 to 0.81) | 0.76 (0.73 to 0.79) | 0.76 (0.72 to 0.79) |

**Table 2.** Prognostic performance (AUROC with 95% confidence intervals) of the models developed in this report and published previously on the combined validation datasets for identification of incipient TB within a 2.5-year period, stratified by mutually exclusive time interval to disease.

| Models | 0-3m to disease | 3-6m to disease | 6-12m to disease | 12-18m to disease | 18-24m to disease | 24-30m to disease |
| --- | --- | --- | --- | --- | --- | --- |
| Full model | 0.94 (0.91 to 0.97) | 0.83 (0.76 to 0.90) | 0.79 (0.72 to 0.86) | 0.76 (0.67 to 0.86) | 0.74 (0.62 to 0.86) | 0.77 (0.57 to 0.98) |
| Reduced model | 0.94 (0.91 to 0.97) | 0.75 (0.67 to 0.83) | 0.84 (0.77 to 0.90) | 0.73 (0.63 to 0.82) | 0.75 (0.64 to 0.87) | 0.89 (0.73 to 1.00) |
| Sweeney 3 | 0.68 (0.63 to 0.74) | 0.44 (0.35 to 0.52) | 0.63 (0.55 to 0.71) | 0.49 (0.39 to 0.58) | 0.47 (0.35 to 0.59) | 0.58 (0.36 to 0.80) |
| RISK 6 | 0.92 (0.89 to 0.95) | 0.75 (0.67 to 0.83) | 0.80 (0.73 to 0.87) | 0.69 (0.60 to 0.79) | 0.67 (0.55 to 0.79) | 0.70 (0.48 to 0.92) |
| BATF2 | 0.68 (0.62 to 0.73) | 0.56 (0.47 to 0.64) | 0.54 (0.47 to 0.62) | 0.50 (0.40 to 0.59) | 0.46 (0.34 to 0.58) | 0.30 (0.14 to 0.47) |
| Suliman 4 | 0.90 (0.86 to 0.93) | 0.70 (0.61 to 0.79) | 0.69 (0.61 to 0.76) | 0.57 (0.47 to 0.67) | 0.62 (0.49 to 0.74) | 0.60 (0.38 to 0.83) |

We tested sensitivities and specificities across models using cut-offs identified by the maximal Youden Index, which is based on the best tradeoff between sensitivity and specificity from each ROC of the time interval. Both of our models met or approximated the minimum criteria (>75% sensitivity and >75% specificity) in the WHO target product profile (TPP) for prediction of progression to TB disease (*50*) over a 30-month period (the full model sensitivity 74.2 % and specificity 78.3%, the reduced model sensitivity 77.3% and specificity 74.9%). RISK6 and Suliman 4 only met the criteria over 0-12 months and 0-6 months, respectively. Sweeney 3 and BATF2 didn’t meet the criteria in any of the time periods (**Supplementary Table 5**). Based on a pre-test probability of 2% with Youden Index cutoff, PPVs and NPVs of the models were estimated. Both of our models achieved a PPV ranging from 16.8% to 17.7% for a 0-3 month and marginally surpassed the WHO target PPV of > 5.8% over 2 years (the full model PPV 6.6% and the reduced model PPV 5.9%). None of the other models except RISK6 met the WHO PPV criteria over 0-2 years (RISK6 PPV 5.9%) (**Supplementary Table 5**). Alternatively, we used a pre-specified cutoff of 2 standard deviations (SDs) above the mean of the non-progressing group to prioritize PPVs and specificity, which was proposed in previous reports (*27, 43*). At this threshold, PPVs estimated in our models significantly improved to 27.9% and 30% over 0-3 months and 19.5% and 20.3% over 0-24 months for the full and reduced model, respectively, which were substantially superior to the WHO target PPV over 0-2 years. Similar improvement of PPVs was also observed in the RISK6 and Suliman 4 models (**Supplementary Table 6**). However, overall sensitivities using this cutoff were significantly reduced compared to the one using Youden Index while specificities were maintained at a high level (> 96% in all models). Sensitivities over 0-3 months are 66.7% for the full model and 69% for the reduced model. Over 0-30 months, sensitivities are 41.1% for both of our models. Many cases at distant time points to disease progression (>12 months) could not be detected by this cutoff.

### Comparison of gene signature models in TB diagnosis

Two published gene signature models (Sweeney 3 and RISK 6) demonstrated their utility as a screening or triage test for TB diagnosis in multicenter prospective studies (*32, 38*). Here we examined diagnostic performance of our models to discriminate active from non-active TB subjects in comparison with the published models, using the pooled datasets of all 37 studies. Using this pooled dataset allowed us to recapitulate various confounding variables to mimic a real-world setting. After pooling the datasets, subjects were stratified by different TB states, and the scores for every subject were generated by individual models. Additionally, as Mulenga *et al*. highlighted, respiratory viral infections induce interferon-stimulated genes, which can be a potential confounding variable for gene signature predictions (*37*). We therefore also included an additional 20 datasets containing 15 different respiratory viral infections (**Supplementary Table 1)**. Note, since our RF models were generated by using part of these pooled datasets (discovery dataset), the assessment of our model performance in this section was subjected to 5-fold nested CV to ensure a blinded model validation. The full model demonstrated score distributions able to distinguish between the groups that provided a strong discrimination between ATB vs. HC, LTBI or OD (**Fig. 3E, Supplementary Fig. 11**). Strikingly, the full model was able to discriminate viral infection (VI) from ATB (AUROC 0.93, 95% CI 0.91-0.94) and overall AUROC between ATB and non-ATB (HC, LTBI, OD and VI) achieved 0.92 (95% CI 0.91-0.93). The reduced model using fewer features reduced the diagnostic performance as expected but maintained an AUROC at 0.87 (95% CI 0.86-0.89) when separating ATB from non-ATB. Its reduced performance was primarily attributed to insufficient differentiation between ATB and OD (AUROC 0.75, 95% CI 0.73-0.77) and between ATB and VI (AUROC 0.83, 95% CI 0.81-0.85) (**Supplementary Fig. 11**). In contrast, the pooled TB scores generated by the published models exhibited multimodal or disperse distributions in multiple groups (**Supplementary Fig. 11**), which resulted in less overall discrimination between ATB and non-ATB (Sweeney 3 AUROC 0.81, 95% CI 0.79-0.83; RISK 6 AUROC 0.75, 95% CI 0.73-0.77; BATF2 AUROC 0.70, 95% CI 0.68-0.72; Suliman 4 AUROC 0.79, 95% CI 0.75-0.82) (**Supplementary Fig. 11**). The results suggest that published models assessing the expression of fewer genes, especially the BATF2 model testing a single gene, may be more sensitive to confounding variables.

### Establishing a probabilistic TB risk assessment model

Identifying a positive case from a screening or triage test for disease diagnosis or prognosis usually requires a pre-defined cutoff, which needs to be validated by post-hoc analyses from independent studies. Given heterogeneous blood transcriptomic responses to TB progression due to confounding variables as previously described, the pre-defined cutoff was often variable in different settings. Similar evidence has been also discussed in other reports (*3, 16, 51*). In addition to using the cutoff approach for test readout, here we also propose a new probabilistic model to estimate TB risk using a cutoff-free method while considering individual and population heterogeneity in the model. Initially, to build the probability model we used our full model to compute all the TB scores and stratified the datapoints into different TB states (**Fig. 4A**). The score distributions were consistently elevated beginning with healthy controls and latent infection without progression, followed by incipient TB groups ranging from distant to proximal to disease. Active TB showed the highest scores. Using stratified score data, we generated a probability model to estimate TB risk as a function of TB scores ranging between 0 to 1. The model was composed of 6 best-fitted probability functions, each of which estimated the probability associated to the observed outcome (healthy control, latent infection without progression, < 24m, < 12m, < 3m to disease, and active TB), for any given TB score (**Fig. 4B**). For instance, a subject with a TB score = 0.2 is estimated to be 82% in LTBI, 58% in HC and only <5% in either incipient or active TB. In contrast, a subject with a TB score = 0.6 is estimated to be 78% in < 24m, 72% in < 12m, 58% in < 3m to disease, 42% in active TB, and only < 5% in either HC or LTBI. As the score increases and approaches 0.8, the probabilities for incipient TB < 3m to disease will dramatically increase and approach 100%, similar to active TB with a slight right shift of the distribution. This probabilistic model allows us to generate an additional readout for TB risk stratification or TB screening while taking into account uncertainty of the data due to biological or technical variation.

**Figure 4.**
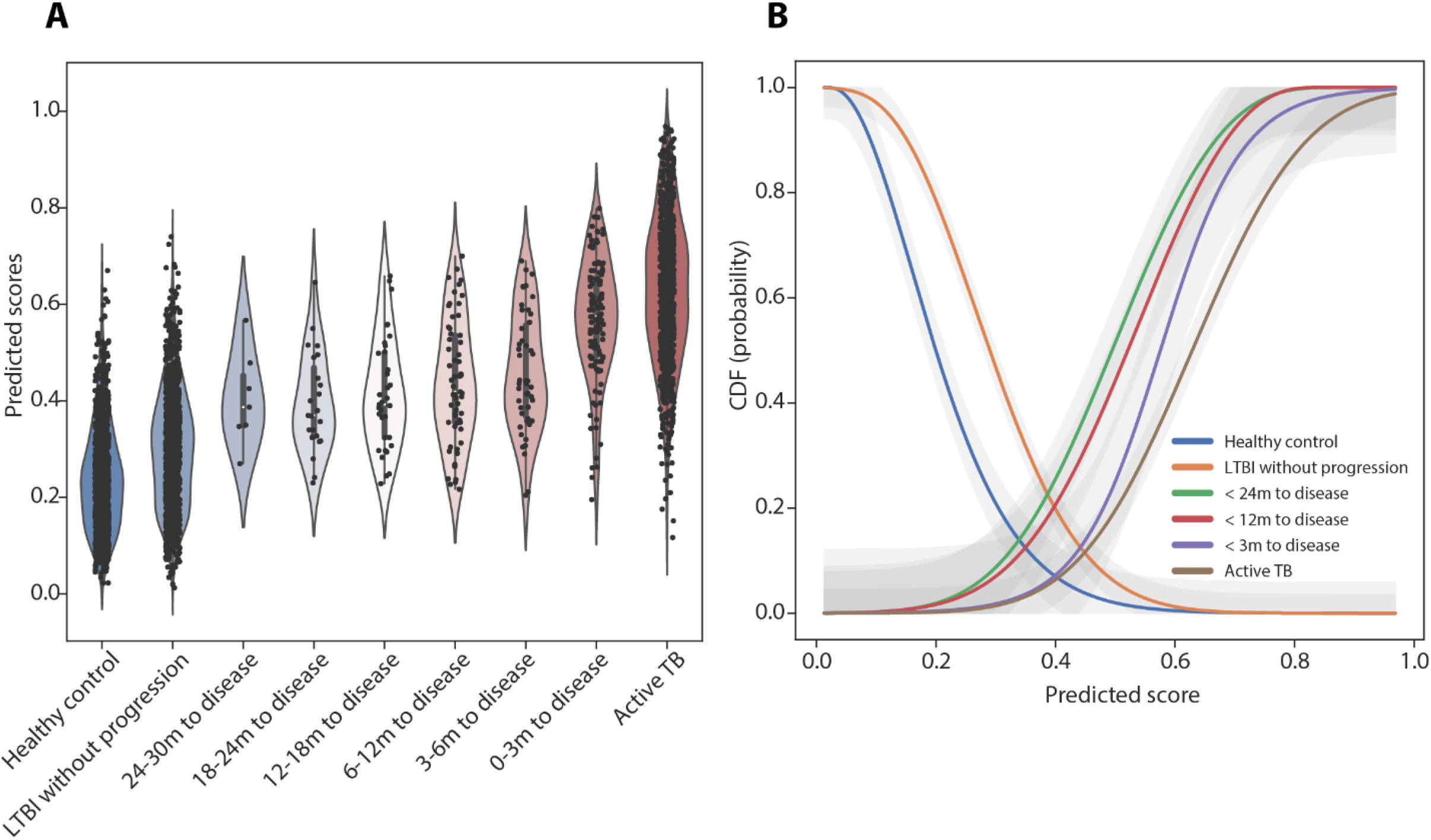
Building a probabilistic TB risk model. (A) The distributions of the full model-generated TB scores, stratified by categorical interval to disease, and active TB, are shown in the violin plot, based on the pooled dataset collected from all the 37 studies. Individual datapoints of the dataset are plotted as a point in the violin plot (datapoints *n* = 3869). (B) The 6 probability functions (cumulative distribution function) that depict the probabilities of the outcomes (HC, LTBI, < 24m, < 12m, < 3m to disease, and active TB) as a function of TB scores were showed in lines with different colors. The shaded area represented the 95% confidence interval area.

### Validation of the gene signature models in treatment monitoring and treatment outcome prediction

We have shown whole blood gene signatures as a correlate of immune status responding to *Mtb* bacterial load during TB progression. Accumulated evidence has also demonstrated the utility of gene signatures in treatment monitoring along with *Mtb* elimination (*3, 16, 30–35*). Here we interrogated whether our gene signature model could be used to infer treatment responses as well as be predictive of clinical outcomes (cure, failure, and recurrence). Two independent treatment studies which provided clinical characterization with publicly available transcriptome and bioassay data were used to assess the consistency of our model prediction with the study results (*31, 35, 52*).

The Catalysis treatment response dataset contained 96 HIV-uninfected adults with newly diagnosed pulmonary TB undergoing standard 6-month treatment (*52*). Of the 96 participants, 8 did not achieve cure and failed treatment when the month 6 sputum culture remained positive, and 12 of 88 cured participants had recurrent TB within 2 years after treatment completion. We first examined the dynamics of TB scores generated by our model in response to treatment. Since the reduced RF model’s performance in the prognostic assessment is non-inferior to the full model (**Tables 1 and 2**), and holds greater potential for clinical development, we solely used the reduced model for the validation in treatment monitoring. The predicted scores significantly declined during the first week after treatment initiation and continuously decreased with treatment duration. At the end of treatment (EOT) (Day 168), the scores from most of the participants were at a similar level as healthy controls (**Fig. 5A**). The model was able to significantly discriminate between healthy controls and before treatment (Day 0) (AUC 0.92, 95% CI 0.87-0.97), 1 week (AUC 0.76, 95% CI 0.67-0.85), and 4 weeks after treatment initiation (AUC 0.72, 95% CI 0.63-0.81). Poor discrimination between healthy controls and EOT was observed which agreed with the clinical outcome where most of the participants (88 out of 96) were cured at EOT (**Fig. 5B**). Interestingly, we further stratified the participants by time to sputum culture conversion to negative (negativity at day 28, 56, 84 and 168, and no conversion at day 168 [treatment failed]). The participants who failed treatment retained high TB scores throughout the course of treatment, which were statistically different from cured groups. In contrast, at 7 days after treatment initiation the TB scores from the group with the earliest culture conversion to negative (day 28) had approached a similar level as at EOT (**Fig. 5C**) (**Supplementary Fig. 12**). Importantly, the reduced model statistically discriminated between participants with bacteriological cure and those with treatment failure not only at EOT (AUC 0.95, 95% CI 0.83-1.00) but at baseline (AUC 0.73, 95% CI 0.51-0.95) (**Fig. 5D**). RISK 6 and BATF2 also provided a comparable result in treatment outcome prediction (**Supplementary Fig. 14A**). Strikingly, when we compared TB scores at baseline from poorly adherent subjects against treatment outcome (patients who missed more than 20% of treatment were stratified in the group of poor adherence (*52*)), the subjects with treatment failure showed statistically higher levels of TB scores than those who were cured (*p* < 0.01) (**Fig. 5E**). While treatment failure was statistically associated with poor adherence in this study (*p* = 0.00032, ANOVA test), we demonstrated TB scores at baseline as another confounding factor impacting treatment outcome. Moreover, with a portion of cured participants in this study developing recurrent disease within 2 years after treatment completion, we investigated whether the gene signature could differentiate the participants with and without recurrence before EOT. The model didn’t provide enough discriminative evidence to separate the two groups at any timepoint during treatment course (**Supplementary Fig. 13**) and neither did the published models (**Supplementary Fig. 14B**). This could be because these individuals were never defined as relapses vs reinfections and thus the models should be retested on validated relapses. Moreover, we investigated if our model contained properties linked to lung inflammation based on [^18^F] FDG PET-CT imaging responses in this study, which were reported in RISK 6 and Sweeney 3 (*3, 44*). The scores generated from our model statistically correlated with not only lung lesion activity measured by total glycolytic ratio activity (TGRA) at three time points (Day 0 Spearman *r* = 0.6, *p* < 0.0001, Day 28 Spearman *r* = 0.66, *p* < 0.0001, and Day 168 Spearman *r* = 0.31, *p* = 0.0022) but also MGIT culture time to positivity (Spearman *r* = −0.62, *p* < 0.0001) and Xpert Ct values (Spearman *r* = − 0.52, *p* < 0.0001) both measured before treatment (**Supplementary Fig. 15A-E**). In addition, the scores at baseline are significantly higher in the participants with radiologically persistent lung inflammation at EOT than those with radiologically cleared lung inflammation (persistent or cleared lung inflammation was defined by the cutoff of TGRA = 400 from the previous report (*52*)(**Supplementary Fig. 15F**). Taken together, these findings suggest that the scores from our model are associated to both metabolic activity of TB lung lesion and active *Mtb* infection, both of which contributed to EOT outcome prediction.

**Figure 5.**
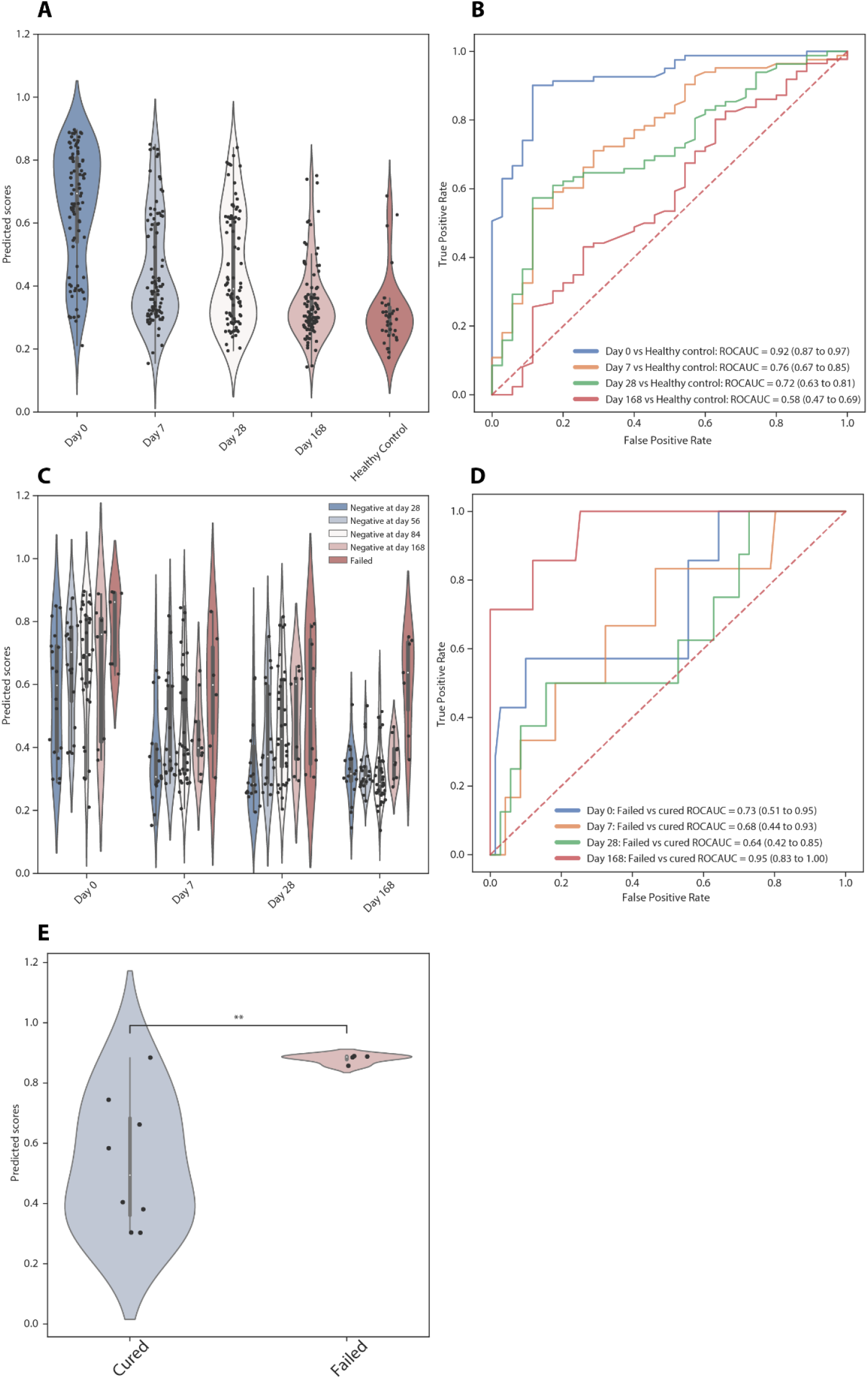
Model validation using the Catalysis TB treatment study. (A) Comparison of TB scores generated by the reduced model at Day 0, 7, 28, and 168 after treatment initiation and from healthy controls are shown in a violin plot. Individual datapoints are plotted as a point in the violin plot (datapoints *n* = 395). (B) Receiver operating characteristic curves depict model discrimination between healthy controls and the patients from different time points after treatment initiation. Area under the curve and 95% confidence intervals for each interval to disease are also shown. (C) At each timepoint the TB scores were stratified by the time of sputum culture conversion to negative (negativity at day 28, 56, 84 and 168, and no conversion at day 168 [failed]) and plotted in the violin plot (datapoints *n* = 360). (D) ROC curves, stratified by different timepoints after treatment initiation, depict predictive performance of the models for discrimination between the patients with bacteriological cure and those with treatment failure at EOT. (E) Comparison of TB scores against treatment outcome from poorly compliant subjects at day 0. P-values were calculated using the Mann-Whitney U test (**, *p* < 0.01).

The Leicester treatment cohort is composed of 74 participants with pulmonary TB. Of these participants,16 received a 6-month standard regimen treatment (clinical cure < 200 days), 39 received extended treatment (requiring >200 days of standard regimen treatment due to clinical suspicion of TB at EOT), 7 were defined as difficult TB cases (requiring extended treatment due to treatment intolerance and/or adherence issues), 4 were TB drug resistance patients, and 8 were infected with a chronic local outbreak TB strain (*31*). Consistent with the Catalysis analysis, declining TB scores after treatment initiation were observed. Despite this, the scores didn’t reach to the same level as healthy control until > 1 year after treatment initiation, which might be explained by 39 out of 74 participants requiring extended treatment (**Fig. 6A**). Consistently, the model significantly discriminated between healthy controls and participants in treatment until month 4. Its performance began to wane during month 4-6 (AUC 0.70, 95% CI 0.61-0.79), and month 7-12 (AUC 0.71, 95% CI 0.63-0.80), at which point a portion of the patients had been cured based on clinical assessment (**Fig. 6B**). The average TB score at baseline was the highest among different time intervals, however the score distribution was scattered with a long tail. Interestingly, we stratified the patients based on smear positive and negative and found that smear positive patients were statistically more likely to have higher TB scores (*p* < 0.0001). Most of the samples in the long tail of the score distribution were from smear negative patients (**Fig. 6C**). Furthermore, the scores from patients requiring extended treatment were consistently higher than those on standard treatment at week 3-4 (*p* > 0.05), month 2-3 (*p* < 0.01) and month 4-6 (*p* < 0.0001) (**Fig. 6D-F**). A previous report identified that smear positive patients mostly fell within the extended treatment patient group (*31*). Our model also showed a consistent prediction where patients requiring extended treatment displayed higher scores throughout the course of treatment (**Fig. 6G**), which suggested that the duration of treatment required in individual patients could be predicted at early timepoints. Indeed, TB scores at month 2-3 were a stronger predictor of treatment duration and significantly discriminated between the patients requiring standard and extended treatment after a 6-month treatment (**Fig. 6H**). In summary, model validation in two independent treatment studies suggests that our model holds potential to monitor TB treatment success and to predict the duration of treatment required for cure at the end of treatment in TB patients.

**Figure 6.**
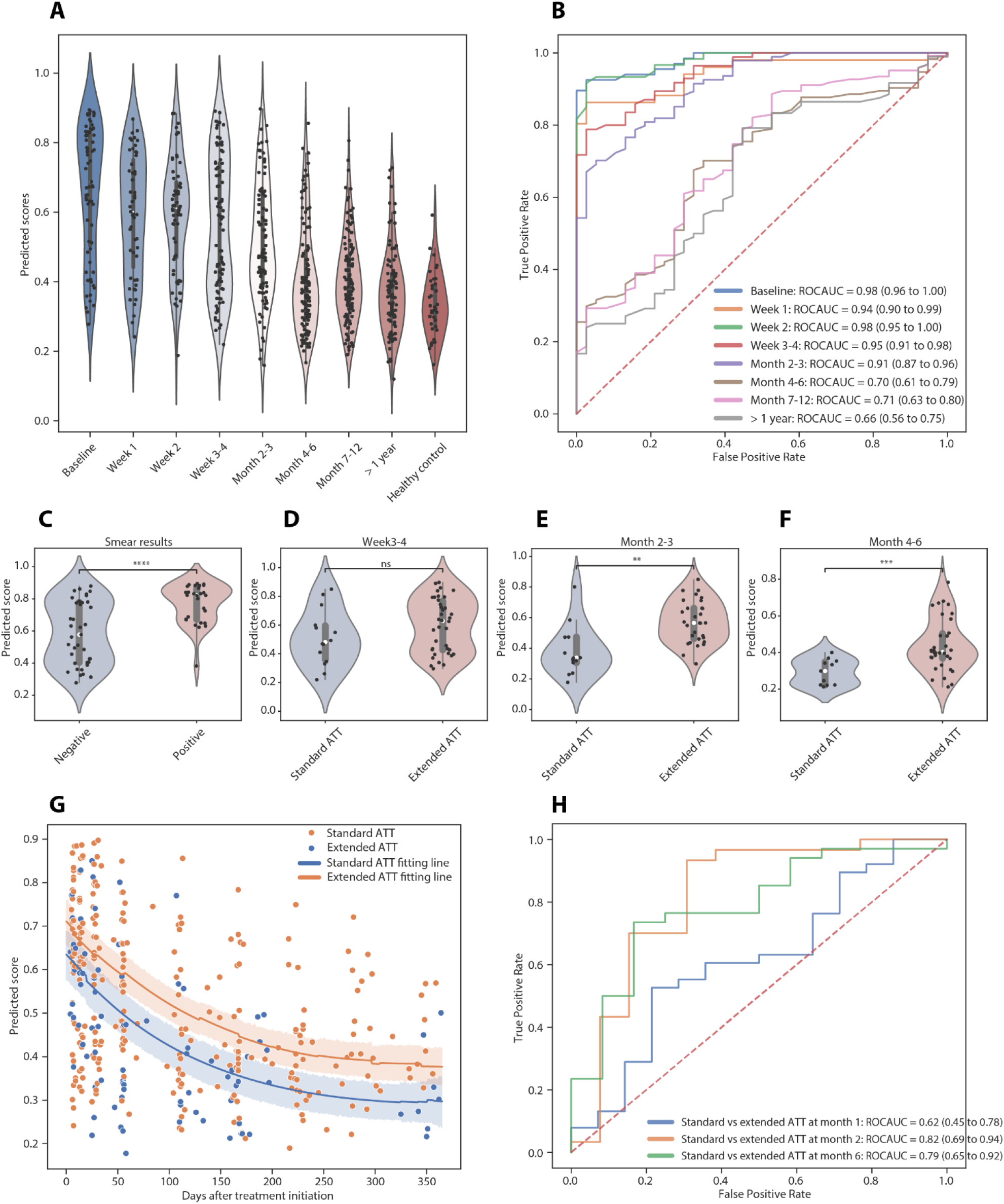
Model validation using the Leicester TB treatment study. (A) The distributions of the reduced model-generated TB scores, stratified by different time intervals after treatment initiation and healthy control, are shown in the violin plot. Individual datapoints are plotted as a point in the violin plot (datapoints *n* = 728). (B) Receiver operating characteristic curves depict model discrimination between healthy controls and the patients from different time intervals after treatment initiation. Area under the curve and 95% confidence intervals for each interval to disease are also shown. Comparison of TB scores between smear positive and negative patients at TB diagnosis (C), between the patients requiring standard and extended anti-TB treatment (ATT) at week 3-4 (D), month 2-3 (E) and month 4-6 (F) after treatment initiation are shown in the violin plots. P-values were calculated using the Mann-Whitney U test (ns, *p* < 1; *, *p* < 0.05; **, *p* < 0.01; ***, *p* < 0.001; ****, *p* < 0.0001). (G) Scatterplots show TB scores throughout treatment course, color-coded by the patients requiring standard or extended ATT. The line represents the median of the stratified group with 95% confidence interval around the median shown in the shaded area. (F) ROC curves, stratified by different timepoints after treatment initiation, depict predictive performance of the models for discrimination between the patients requiring standard and extended ATT.

## DISCUSSION

Here we developed a network-based meta-analysis with ML modeling to generate a common 45-gene signature specific to active TB disease and to systematically train/validate a generalized predictive model using 57 studies with 6290 data points in total. By leveraging heterogeneity among the included cohorts and implementing rigorous feature selection and model hyperparameter optimization, we demonstrated robust performance of the model in both short-term and long-term TB risk estimation and treatment monitoring that provides a complementary approach to the current models, most of which offer good performance in ATB diagnosis and/or short-term TB risk estimation. Importantly, our model also showed robust discrimination between ATB and different respiratory viral infections, demonstrating its predictive specificity to ATB. None of the previous models have yet been systematically validated in this aspect.

As many TB gene signatures have been reported previously, we collected 30 previously published gene signatures (**Supplementary Fig. 5**) (*2, 43*), and found that a large portion of the genes (563 out of total unique 721 genes) were only detected once. This limited commonality among the signatures could be driven by heterogeneity in cohorts or different computational approaches. In comparison of our 45-gene set with 30 previously published gene signatures, 25 out of 45 genes were included in at least three published gene signatures, and particularly genes GBP5, C1QB, FCGR1B, DUSP3, and BATF2 have repeatedly identified in many reports (**Supplementary Fig. 5**), suggesting the reproducibility of our network-based meta-analysis approach to generate a consistent result. The remining 20 genes, regardless of having only one match or no match to other gene signatures, demonstrates an association to active TB (**Fig. 2E-H**), may potentially play a role in increasing model sensitivity for long-term TB risk estimation and improving model discrimination between TB disease and viral infection. Along this line, we demonstrated the 45-gene set responding consistently to active TB across cohorts versus other conditions (HC, LTBI, and OD) (**Fig. 2E-H**). More importantly, we also observed that the subjects with HIV and TB coinfection exhibit a similar expression pattern in the 45-gene set when we stratified the active TB subjects into HIV positive and negative in the heatmap (**Fig. 2E-H**). This provides additional evidence that our gene signature can be applied for TB risk estimation in high HIV burden countries.

While building a predictive model to maximize the orthogonal information of the 45 common genes that is associated to the cohort outcomes, we developed three constitutive steps – taking the ratio of the paired gene expressions, feature down-selection, and supervised machine learning in pooled heterogenous cohorts. Feeding the paired gene ratio into the predictive model not only considers first-order interaction between the genes, but also allows accommodation of various data scales generated by different profiling technologies. To produce a generalizable supervised learning model, we maintained cohort-to-cohort variations by not performing data correction on confounding variables. In addition, the model was regressed to two states: ATB and non-ATB containing HC, LTBI, OD or Tx, which enforces the model maximizing the variation between ATB and non-ATB while considering the difference among the conditions within non-ATB. Finally, ML model benchmarking (**Supplementary Fig. 6, Supplementary Table 3**) and its hyperparameter optimization with a framework of repeated nested cross validation allow us to identify a ML model to be generalizable for independent cohorts. While generating the reduced model, we implemented a feature stability selection approach, a sensitivity analysis to interrogate the likelihood of a feature selected in the model based on randomly generating subsets of a dataset. The list of feature rankings, which refers to the importance of the features contributing to model prediction is displayed in **Supplementary Fig. 7B**. The reduced model is currently considered to use the minimal number of the features that produced sufficient discriminative power as compared to the full model (**Supplementary Fig. 8)**. When applying for a PoC screening test, other factors including selection of the technology either qRT-PCR (*38*) or NanoString (*53*) and the cost of the assay will also need to be considered. Using the feature ranking list as a guidance, we can generate a new model by adding or removing genes based on the needs in the PoC setting. However, to maintain the maximum value to the patient it might be desirable that technology improve to utilize the signature rather than the signature be reduced to accommodate the technology.

In comparison of model performance in TB prognosis and diagnosis **(Fig. 3, Supplementary Fig. 9-11)**, we included four published models in this report, each of which has reduced the number of the genes required to the minimum in order to develop a PoC screening or triage test. Most of the genes selected in the models are strongly associated with active TB (*3, 5, 18, 27*). However, when the models were tested in the pooled dataset compiled from multiple cohorts, we observed that the TB scores particularly from non-active TB groups, such as HC and LTBI, showed much larger dispersion in the distributions than from the active TB group (**Supplementary Fig. 10-11)**. For instance, the scores from Sweeney 3 in the group of LTBI without progression in 2 years exhibited a clear bimodal distribution due to a combination of two different TB score levels generated from the different cohorts (**Supplementary Fig. 10A)**. As a result, we observed weak discrimination between the non-progressing and progressing groups. BATF2, a single gene model, demonstrated accurate discrimination between active and latent TB (*20, 27*), however, showed low discriminatory power in a multi-cohort setting with different confounding variables **(Supplementary Fig. 11I-J)** (similar cases in other models can be also found in **Supplementary Fig. 11**). This collective evidence via a meta-analysis suggests that the minimized gene signatures may respond not only to active TB disease, but to other diseases or physiological conditions. As Mulenga et al highlighted (*37*), individuals with respiratory viral infection had elevated RISK 11 gene signature scores which made it difficult to differentiate between active TB and viral infection. Our full model showed less dispersed distributions in non-active TB groups (**Fig. 3A, Supplementary Fig. 11A-B**) and provided strong discrimination between ATB and other groups, including viral infection. Our reduced model using a part of the full model gene signature maintained good discrimination between ATB and non-ATB, but its performance was reduced against viral infection and other lung diseases (**Supplementary Fig. 11 C-D**). Taken together, the results highlight that model only considering the minimized gene set (the best predictors) may not be sufficient to deal with cohort heterogeneity. A model containing a combination of genes with both strong and moderate predictors may improve its reliability in the context of heterogeneity and gain model specificity to TB.

From TB treatment studies, our gene signature model demonstrates that TB score dynamics nicely correlate with treatment duration (**Fig. 5A and 6A**), although individual variations are noticeable (**Supplementary Fig. 12**), which might result from varied pathophysiological conditions at baseline (*54*). In addition, the values of the TB scores in our model are associated with lung inflammation during treatment (**Supplementary Fig. 15**). This suggests that the TB scores as a biomarker of host immune status respond to *Mtb* bacterial load and its changes ongoing in the lungs. Medication adherence is one of the confounding factors to evaluate treatment outcomes (cure and failure). In the Catalysis study, by dissecting different confounding factors we demonstrated that both drug adherence and *Mtb* bacterial load at baseline, captured by TB score, were individually associated to treatment outcomes (**Fig. 5D-E)**. It’s possible that the signature that predicted treatment failure for these poorly adherent patients-before treatment was initiated - was able to identify a metabolic state that correlated to high bacterial load and thus a likelihood of poor outcome. This has been observed again in a study specifically aimed at metabolic differences of TB patients (*55*). As a caveat, the sample size for the subjects with poor adherence and also for the treatment failure is small and requires additional studies to validate the result.

We demonstrated a complete model development from collecting massive study transcriptomic datasets, identifying a common TB gene signature, generating a generalized predictive model, and systematically validating the model in a real-world setting. This work paves the way for us to further investigate the relationship between TB score and clinical characteristics, outcomes, and other biomarker measurements in our TB treatment clinical trial studies, and to refine our predictive modeling work with other biomarkers to support future clinical trial development and patient care in general.

## MATERIALS AND METHODS

### Transcriptome datasets

We collected 37 publicly available transcriptome datasets from NCBI GEO and EMBL ArrayExpress from previously published studies (**Supplementary Table 1**). Among the datasets, there were 27 studies containing one or few clinical TB states including HC, LTBI, ATB, and treatment (Tx) or other lung diseases (OD) (datapoints *n* = 2914), and 10 progression or treatment studies with longitudinal timepoints (datapoints *n* = 1281). Of these, 26/37 datasets were derived from microarray experiments and 11/37 were derived via RNAseq. We used 27 out of 37 studies (most are the cross-sectional studies) as the discovery datasets to build the networks for our common gene signature discovery, and to train/test the ML model. The remaining 10 longitudinal cohorts as the validation datasets were used to independently validate prediction performance of the ML model on TB prognosis, diagnosis, and responses to TB drug treatment. To further test model specificity to TB disease, we collected additional 20 microarray datasets from GEO (datapoints *n* = 2095) from patients across 12 countries with multiple respiratory viral infections (**Supplementary Table 1**).

### Processing of microarray datasets

Microarray data were downloaded from GEO using the function `getGEOData` from the MetaIntegrator package v2.1.3 with option `qNorm=FALSE` (*56*). Missing values in each expression matrix (probes x individuals) were replaced with 0 before performing quantile normalization with the package preprocessCore v1.55.2. Rows were reduced for each dataset by retaining only a single probe for each gene (the probe with the highest gene expression sum across all samples in the matrix). This transformed and filtered expression matrix was used for downstream analysis.

### Processing of RNAseq datasets

RNAseq raw data were downloaded from NCBI Sequence Read Archive (SRA) and processed through our customized bulk RNAseq pipeline. Fastp v0.20.1 (*57*) with option `qualified_quality_phred = 20` was used to perform quality control, adapter/tail trimming and read filtering on FASTQ data, followed by Salmon v1.4.0 (*58*) with options `recoverOrphans = true, gcBias = true`, and `seqBias = true` for quantification of gene transcript expression from the sequence reads. The trimmed and filtered reads were mapped to the human genome Ensembl GRCh38 (release 102). A read counts matrix was generated from the pipeline for downstream analysis.

### Differential gene expression analysis

We performed pairwise differential gene expression analysis on the 27 discovery datasets used in the network analysis. To run differential gene expression between two groups within a given dataset, we required that at least three individuals belong to each group. Four pairwise comparisons were considered for each dataset: (1) ATB vs. HC, (2) ATB vs. LTBI, (3) ATB vs. OD, and (4) ATB vs. Tx.

#### Microarray Data

We used the limma package v3.46.0 to create the contrasts matrix (using functions `lmFit` and `makeConstrasts`) and fit a linear model for each probe (gene)(*59, 60*). Finally, we used the functions `eBayes` with option `proportion = 0.01` and `topTable` with option `adjust = “fdr”` to compute the log-odds of differential expression for each gene and correct the p-values for multiple hypothesis testing by the Benjamini-Hochberg (BH) method.

#### RNAseq Data

We used `calcNormFactors` from the edgeR package v3.32.1 to calculate the normalization factors, then filtered out lowly-expressed genes from each matrix (retaining genes which had at least 1 count-per-million in at least N/1.5 individuals, where N is the number of individuals) (*61*). We then used `voom` from the limma package to transform our dataset before creating the contrast matrix. The remaining steps for linear modeling and expression variability estimation are described previously in microarray data.

### Building gene covariation networks

We constructed four gene covariation networks corresponding to four pairwise comparisons as follows: (1) ATB vs. HC, (2) ATB vs. LTBI, (3) ATB vs. OD, and (4) ATB vs. Tx.

For each comparison, we collected ***M*** datasets for each of which we ran a differential gene expression analysis between the groups. From the results, we retained only genes that had both, (1) a corresponding false discovery rate of 5% or less (BH adjusted p-value < 0.05) and (2) an effect size > 1.5-fold (*logFC* > *log*_2_(1.5)). For each dataset, we then constructed a 1 × ***N*** vector that presented ***N*** genes and contained the *logFC* for each gene that passed the above filters or 0 otherwise. We combined vectors from ***M*** datasets to generate an ***N*** × ***M*** matrix and convert the matrix to +1 if the corresponding *logFC* > 0, 0 if the corresponding *logFC* = 0, or −1 if the corresponding *logFC* < 0 to create matrix ***X*** which held information corresponding to which genes were significantly upregulated and downregulated across all datasets (**Fig. 1A**).

We constructed a covariation network using NetworkX v2.6.3 (https://networkx.org/) with ***N*** nodes (each node represents a gene). Edges between nodes were weighted by considering the covariation between each pair of genes in ***X***. We calculated the dot product between each pair of genes (a pair of rows in ***X***) to generate a distribution (***D***) of edge weights (range - ***M*** to ***M, M*** = number of datasets). To filter out low-value non-zero edges that may have occurred by chance, we performed a permutation test on ***X***. We constructed a null distribution of edge weights by randomly shuffling the columns of ***X*** within each row (**Supplementary Fig. 1**). We repeated this process twenty-five times to create the null distribution (***D’***). For each distribution, ***D*** and ***D’***, we computed the proportion of edge weights to the left of each edge weight for *t* ∈ {-***M***, 0} (normalized by all negative edge weights) and the proportion of edge weights to the right of each edge weight for *t* ∈ {0, ***M***} (normalized by all positive edge weights). Let ***t*** be the proportion of edge weights computed from ***D*** and ***t’*** the proportion of edge weights computed from ***D’***. A threshold ***T***_***l***_ for edge weights < 0 and ***T***_***r***_ for edge weights > 0 was determined by the exact *p*-value 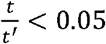. The edges within ***T***_***l***_ and ***T***_***r***_ were then excluded in the network.

### Identifying common genes among the networks

First, the weighted degree of a node, the sum of the edge weights of a node, was calculated to represent the degree centrality of a gene in the network (**Supplementary Fig. 2**). A gene with higher degree centrality indicates its response to ATB more consistently across the cohorts. To look for the most common genes in the network, we modeled the weighted degree distribution based on the assumption that the covariation network is a scale-free network (*62*) where the degree distribution follows a power law (**Supplementary Fig. 3**). Therefore, a power-law distribution (a generalized beta distribution) is used to estimate the weighted degree distribution ***W*** and the cumulative distribution function (CDF) of ***W*** was also estimated. For each network, we identified the genes within the top 5 % of ***W***. Then we collected the top-ranked genes that occurred in at least two of the networks. Next, we excluded genes with *abs*(*mean*(*logFC*)) < 0.5. The final common gene list was composed of 45 genes, where 41 genes were up-regulated and 4 genes were down-regulated in ATB relative to other disease conditions.

### Pathway enrichment analysis

To conduct pathway enrichment, we used REACTOME pathway modules taken from MSigDB (https://www.gsea-msigdb.org/gsea/msigdb/). We ran our 45-candidate gene set through Enrichr (*63*) as implemented in Python through the package GSEApy v0.10.1 (https://github.com/zqfang/GSEApy).

### Establishing optimized predictive models

#### Data preparation and harmonization

Before compiling all the discovery datasets, the data were transformed to logarithmic scale and standardized by z-score within each dataset. The compiled matrix was filtered to contain the common 45 gene expression profiles from each cohort. The outcomes (TB disease states) of the cohorts were united into two distinct states from model building: 1 (ATB) and 0 (HC, LTBI, Tx or OD)

#### Feature generation and selection

To build a predictive model able to handle data generated across different platforms (RNAseq, microarray, or qRT-PCR) without further data transformation, an expression ratio between a pair of genes was used as a feature importing into the model. By considering all pairs of 45 common genes, a 990-dimensional feature vector was constructed. The dimensionality of the feature space was reduced by selecting the discriminant features to obtain a more generalizable and more accurate model. Therefore, two-step feature selection was placed in our framework. 1) Univariate feature selection: mutual information is calculated to measure the dependency between individual feature and the outcome. The top 90% of total features were maintained in the feature space. 2) Multivariate feature selection. We implemented two approaches in our modeling framework for different purposes. LassoLars is a LASSO model, a *l*1-penalized regression model, which generates sparse estimators and whose parameters are estimated by the LARS algorithm (*64*). This approach was considered when a predictive model with a full set of the selected features was generated to achieve the best prediction performance. The second approach was LASSO with stability selection (*65*), where subsampling from the data in conjunction with *l*1-penalized estimation was implemented to estimate the probability for each feature to be selected. For each feature (*k*), the stability path is generated by the selection probabilities 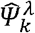 as a function of the regularization parameter (*λ*) when randomly resampling from the data, and the probability (***P****\**) of the feature was determined by 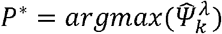 (**Supplementary Fig. 7)**. The features were then ranked by the probabilities and down-selected using a pre-defined cutoff (**Supplementary Fig. 8)**. This approach was considered when only a small number of features was allowed for qRT-PCR to adapt for ultimate translation to a PoC platform.

#### Machine-learning model optimization

We aimed to construct a supervised learning model that took the common gene signature and produced a disease score ranged between 0 and 1 to recapitulate the spectrum of TB infection to disease as well as responses to treatment. To do so, seven state-of-the-art ML regression algorithms in scikit-learn v1.0.1 ML library were tested, including random forest, elastic-net, support vector machine, adaptive boosting, partial least squares, multi-layer perceptron, and extreme gradient boosting.

To establish an optimized model without overfitting, a framework of 5-repeated 5-fold nested cross-validations (CV) was designed (*66*). In each repetition, the dataset was randomly divided into 5 folds, where 80% of the dataset as the training set was used for model building (inner CV) and the remaining hold-out dataset as the test set was used to estimate the true prediction error of the model (outer CV), where the goodness-of-fitness of the model was measured by mean squared error (MSE) and R-squared between the predicted outcome and the actual. Of note, due to the unbalanced outcomes (TB states 0 or 1) in the dataset, each fold was stratified to maintain the similar proportion of each state as the complete dataset. In inner CV, the model was trained the following way. First, the bottom 10 % features were filtered out by using univariate feature selection (as described above) with the whole training set. Secondly, the training set was divided again based on the 5-fold CV strategy. The feature set was further refined using either one of multivariate feature selection approaches (as described above) based on minimization of CV error estimates. Third, to pinpoint the optimal hyperparameters of the learning algorithm, a randomized search in high-dimensional parameter spaces (*67*) was applied to increase search efficiency as compared to the brute-force search approach, and the final parameter setting was determined by the one generating the minimal CV errors. Finally, the true prediction error of the model generated from the inner CV was estimated using the test set in the outer CV. To benchmark the regression algorithms, both the area under the receiver operating characteristic curve (AUROC) and MSE were used to assess the true error of the model (**Supplementary Fig. 6, Supplementary Table 3**).

After the model and its optimal parameters was determined in 5-repeated 5-fold nested CV, the final model was re-trained using the whole discovery dataset (27 cohorts) and its actual prediction performance in progression and responses to treatment were systematically validated by the longitudinal validation dataset (10 cohorts) and viral infection datasets (20 cohorts). In the end, we finalized two models based on different multivariate feature selection approaches – the full model with full-set selected features that maximize prediction performance and the reduced model with reduced features that retains comparable performance (**Supplementary Table 4**).

### Model validation in TB prognosis and responses to treatment

For the progression datasets (7/10 validation cohorts), the datasets were directly compiled without further batch correction to mimic the real-world situation where the prognostic triage tool is performed at an individual subject level. AUC ROC was assessed to evaluate predictive performance that discriminated between the progression group in different prespecified intervals to disease (< 3 months, < 6 months, < 12 months, < 18 months, < 24 months, and < 30 months) and the group including LTBI without progression during the 2-year follow-up. Additionally, sensitivity, specificity, and positive/negative predictive values (PPVs/NPVs) were assessed for each of these time intervals, when assuming 2% pre-test probability, based on two parameters - 1) the predetermined cutoffs defined by 2 standard deviations (SDs) above the mean of the control group(*27*) to prioritize PPVs and specificity, 2) the maximal Youden Index from AUC ROC which provides the best tradeoff between sensitivity and specificity. Furthermore, to determine the uncertainty of the estimates of model performance metrics, 95% confidence intervals (CIs) were calculated as described in Ying et al (*68*). To investigate the changes of the model-predicted TB scores along with disease progression among the cohorts, the data points were binned using mutually exclusive time intervals to disease of 0-3 months, 3-6 months, 6-12 months, 12-18 months, 18-24 months, 24-30 months, and >30 months. For the treatment cohorts (3/10 validation cohorts), AUC ROC was assessed to evaluate the performance of treatment monitoring that distinguished active TB at baseline and at timepoints after treatment initiation. Furthermore, we also examined the prediction of treatment outcomes (cure, failure, and recurrence) using the data at early timepoints (baseline, week 1, week 2, or month 1), assessed by AUC ROC. To study TB score dynamics over the treatment course across cohorts with different sample collection schedules, the data points were bucketed into several representative time points.

### Developing a probabilistic TB risk model

To accommodate the heterogeneity of immune responses to TB progression in different demographic groups, a probabilistic model was developed to incorporate the uncertainty of the responses associated with the clinical outcomes. To do so, we first compiled all the transcriptomic data (37 cohorts) and extracted the data that was on the trajectory of TB progression as well as from HC, LTBI, and ATB. TB scores for each sample were generated using our predictive model. Next, we focused on six groups – HC, LTBI, < 24 month to disease, < 12 month to disease, < 3 month to disease, and ATB. For each group, we modeled TB scores using a best-fitted probability distribution in the following steps. Initially, the data was to fit to various distributions in which SciPy v1.7.3, a Python library, has a list of predefined continuous distributions that can be tested against the data, and the best-fitted distribution with appropriate parameters estimated was identified by using sum of square error between the actual and fitted distributions. Next, given the best-fitted distribution, its probability density function (PDF) and cumulative density function (CDF) were estimated. Therefore, consider TB score as a continuous random variable ***X*** with PDF *f*_*X*_ (*x*), the CDF can be obtained from:

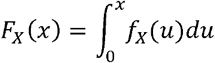

Since TB score is ranged between 0 and 1, the area under PDF between 0 and 1 must be one.

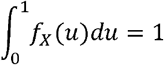

For the groups of different time intervals to disease and ATB, we calculated the cumulative probability that TB score takes a value less than or equal to *t*:

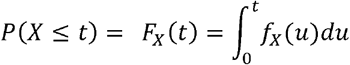

Conversely, for the groups of HC and LTBI, we calculated the cumulative probability that TB score takes a value large than or equal to *t*:

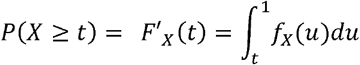

The cumulative probabilities will be used to estimate the probability associated to each of six groups, given a TB score from individuals. To determine 95 % CIs of CDF, the Dvoretzky-Kiefer-Wolfowitz inequality was used to generate a CDF-based confidence band (*69*).

### Statistics

A two-sided Wilcoxon Rank Sum Test was used for pairwise analyses comparing TB disease scores across any clinically defined groups. The BH method was used to calculate false discovery rate (FDR)-adjusted *p*-values for multiple-comparison post-hoc correction. An adjusted *p*-value of less than 0.05 was considered significant. Spearman’s rank analysis was used to test for correlations between variables.

### Code

For RNAseq processing pipeline, we used Nextflow (https://www.nextflow.io/) for analytics workflow development and workflow submission into Slurm, a workload manger, to distribute computing jobs on AWS HPC parallel cluster. Microarray/RNAseq data process and differential gene expression analysis were conducted in R with customized scripts. The remaining works, including network analysis, supervised learning model building and validation, and probabilistic modeling were then established in Python. The code for machine learning was adapted to the multi-processing framework to allow parallel computing by leveraging AWS EC2 instance with 64 CPUs.

### Data and code availability

All data used in this report were collected from publicly available databases (NCBI GEO and EMBL ArrayExpress) and the list of the accession ID of the datasets are available on supplementary table 1. All the code for statistical modeling and figure visualization can be accessed on GitHub (https://github.com/wenhan-yu/tb-common-gene-signature).

## ACKNOWLEDGEMENTS

The authors appreciate Dr. Jill Winter for concept discussion and general feedback in the manuscript, Justin Zhang for viral infection dataset collection, Dr. Aparna Anderson for feedback in statistical modeling, and Dr. Gerard Tromp for feedback in bioinformatics approaches.

## Notes

### Competing Interest Statement

The authors have declared no competing interest.

### Summary of Updates

Correct author affiliation

## REFERENCES

1. World Health Organization, Global tuberculosis report (2021).

2. H. Warsinske, R. Vashisht, P. Khatri, Host-response-based gene signatures for tuberculosis diagnosis: A systematic comparison of 16 signatures. PLoS Med 16, e1002786 (2019).

3. A. Penn-Nicholson et al., RISK6, a 6-gene transcriptomic signature of TB disease risk, diagnosis and treatment response. Sci Rep 10, 8629 (2020).

4. T. J. Scriba et al., Biomarker-guided tuberculosis preventive therapy (CORTIS): a randomised controlled trial. The Lancet Infectious Diseases 21, 354–365 (2021).

5. T. E. Sweeney, L. Braviak, C. M. Tato, P. Khatri, Genome-wide expression for diagnosis of pulmonary tuberculosis: a multicohort analysis. Lancet Respir Med 4, 213–224 (2016).

6. S. T. Anderson et al., Diagnosis of childhood tuberculosis and host RNA expression in Africa. N Engl J Med 370, 1712–1723 (2014).

7. M. P. Berry et al., An interferon-inducible neutrophil-driven blood transcriptional signature in human tuberculosis. Nature 466, 973–977 (2010).

8. C. I. Bloom et al., Transcriptional blood signatures distinguish pulmonary tuberculosis, pulmonary sarcoidosis, pneumonias and lung cancers. PLoS One 8, e70630 (2013).

9. L. Laux da Costa et al., A real-time PCR signature to discriminate between tuberculosis and other pulmonary diseases. Tuberculosis (Edinb) 95, 421–425 (2015).

10. M. Jacobsen et al., Candidate biomarkers for discrimination between infection and disease caused by Mycobacterium tuberculosis. J Mol Med (Berl) 85, 613–621 (2007).

11. M. Kaforou et al., Detection of tuberculosis in HIV-infected and -uninfected African adults using whole blood RNA expression signatures: a case-control study. PLoS Med 10, e1001538 (2013).

12. S. Leong et al., Existing blood transcriptional classifiers accurately discriminate active tuberculosis from latent infection in individuals from south India. Tuberculosis (Edinb) 109, 41–51 (2018).

13. J. Maertzdorf et al., Concise gene signature for point-of-care classification of tuberculosis. EMBO Mol Med 8, 86–95 (2016).

14. A. Sambarey et al., Unbiased Identification of Blood-based Biomarkers for Pulmonary Tuberculosis by Modeling and Mining Molecular Interaction Networks. EBioMedicine 15, 112–126 (2017).

15. L. M. Verhagen et al., A predictive signature gene set for discriminating active from latent tuberculosis in Warao Amerindian children. BMC Genomics 14, 74 (2013).

16. F. Darboe et al., Detection of Tuberculosis Recurrence, Diagnosis and Treatment Response by a Blood Transcriptomic Risk Signature in HIV-Infected Persons on Antiretroviral Therapy. Front Microbiol 10, 1441 (2019).

17. D. E. Zak et al., A blood RNA signature for tuberculosis disease risk: a prospective cohort study. The Lancet 387, 2312–2322 (2016).

18. S. Suliman et al., Four-gene Pan-African Blood Signature Predicts Progression to Tuberculosis. Am J Respir Crit Care Med, (2018).

19. S. Leong et al., Cross-validation of existing signatures and derivation of a novel 29-gene transcriptomic signature predictive of progression to TB in a Brazilian cohort of household contacts of pulmonary TB. Tuberculosis (Edinb) 120, 101898 (2020).

20. J. K. Roe et al., Blood transcriptomic diagnosis of pulmonary and extrapulmonary tuberculosis. JCI Insight 1, e87238 (2016).

21. J. E. Gjoen et al., Novel transcriptional signatures for sputum-independent diagnostics of tuberculosis in children. Sci Rep 7, 5839 (2017).

22. H. D. Gliddon et al., Identification of Reduced Host Transcriptomic Signatures for Tuberculosis Disease and Digital PCR-Based Validation and Quantification. Front Immunol 12, 637164 (2021).

23. H. H. Huang, X. Y. Liu, Y. Liang, H. Chai, L. Y. Xia, Identification of 13 blood-based gene expression signatures to accurately distinguish tuberculosis from other pulmonary diseases and healthy controls. Biomed Mater Eng 26 Suppl 1, S1837–1843 (2015).

24. L. S. de Araujo et al., Transcriptomic Biomarkers for Tuberculosis: Evaluation of DOCK9. EPHA4, and NPC2 mRNA Expression in Peripheral Blood. Front Microbiol 7, 1586 (2016).

25. Z. Qian et al., Expression of nuclear factor, erythroid 2-like 2-mediated genes differentiates tuberculosis. Tuberculosis (Edinb) 99, 56–62 (2016).

26. J. V. Rajan et al., A Novel, 5-Transcript, Whole-blood Gene-expression Signature for Tuberculosis Screening Among People Living With Human Immunodeficiency Virus. Clin Infect Dis 69, 77–83 (2019).

27. J. Roe et al., Blood Transcriptomic Stratification of Short-term Risk in Contacts of Tuberculosis. Clin Infect Dis 70, 731–737 (2020).

28. A. Singhania et al., A modular transcriptional signature identifies phenotypic heterogeneity of human tuberculosis infection. Nat Commun 9, 2308 (2018).

29. N. D. Walter et al., Blood Transcriptional Biomarkers for Active Tuberculosis among Patients in the United States: a Case-Control Study with Systematic Cross-Classifier Evaluation. J Clin Microbiol 54, 274–282 (2016).

30. A. J. Zimmer et al., A novel blood-based assay for treatment monitoring of tuberculosis. BMC Res Notes 14, 247 (2021).

31. O. Tabone et al., Blood transcriptomics reveal the evolution and resolution of the immune response in tuberculosis. J Exp Med 218, (2021).

32. R. Bayaa et al., Multi-country evaluation of RISK6, a 6-gene blood transcriptomic signature, for tuberculosis diagnosis and treatment monitoring. Sci Rep 11, 13646 (2021).

33. J. Heyckendorf et al., Prediction of anti-tuberculosis treatment duration based on a 22-gene transcriptomic model. Eur Respir J, (2021).

34. N. P. Long et al., A 10-gene biosignature of tuberculosis treatment monitoring and treatment outcome prediction. Tuberculosis (Edinb) 131, 102138 (2021).

35. E. G. Thompson et al., Host blood RNA signatures predict the outcome of tuberculosis treatment. Tuberculosis (Edinb) 107, 48–58 (2017).

36. P. K. Drain et al., Incipient and Subclinical Tuberculosis: a Clinical Review of Early Stages and Progression of Infection. Clin Microbiol Rev 31, (2018).

37. H. Mulenga et al., Longitudinal Dynamics of a Blood Transcriptomic Signature of Tuberculosis. Am J Respir Crit Care Med, (2021).

38. J. S. Sutherland et al., Diagnostic accuracy of the Cepheid 3-gene host response fingerstick blood test in a prospective, multi-site study: interim results. Clin Infect Dis, (2021).

39. D. Szklarczyk et al., STRING v11: protein–protein association networks with increased coverage, supporting functional discovery in genome-wide experimental datasets. Nucleic Acids Research 47, D607–D613 (2018).

40. F. Yi et al., Transcriptional Profiling of Human Peripheral Blood Mononuclear Cells Stimulated by Mycobacterium tuberculosis PPE57 Identifies Characteristic Genes Associated With Type I Interferon Signaling. Front Cell Infect Microbiol 11, 716809 (2021).

41. K. B. Nguyen et al., Interferon alpha/beta-mediated inhibition and promotion of interferon gamma: STAT1 resolves a paradox. Nat Immunol 1, 70–76 (2000).

42. M. L. Donovan, T. E. Schultz, T. J. Duke, A. Blumenthal, Type I Interferons in the Pathogenesis of Tuberculosis: Molecular Drivers and Immunological Consequences. Front Immunol 8, 1633 (2017).

43. R. K. Gupta et al., Concise whole blood transcriptional signatures for incipient tuberculosis: a systematic review and patient-level pooled meta-analysis. The Lancet Respiratory Medicine 8, 395–406 (2020).

44. H. C. Warsinske et al., Assessment of Validity of a Blood-Based 3-Gene Signature Score for Progression and Diagnosis of Tuberculosis, Disease Severity, and Treatment Response. JAMA Netw Open 1, e183779 (2018).

45. P. K. W. Kwan et al., A blood RNA transcript signature for TB exposure in household contacts. BMC Infect Dis 20, 403 (2020).

46. S. Y. Bah, T. Forster, P. Dickinson, B. Kampmann, P. Ghazal, Meta-Analysis Identification of Highly Robust and Differential Immune-Metabolic Signatures of Systemic Host Response to Acute and Latent Tuberculosis in Children and Adults. Front Genet 9, 457 (2018).

47. M. Zhang et al., CD19(+)CD1d(+)CD5(+) B cell frequencies are increased in patients with tuberculosis and suppress Th17 responses. Cell Immunol 274, 89–97 (2012).

48. N. Ahn et al., The Interferon-Inducible Proteoglycan Testican-2/SPOCK2 Functions as a Protective Barrier against Virus Infection of Lung Epithelial Cells. Journal of virology 93, e00662–00619 (2019).

49. C. G. Weindel et al., LRRK2 maintains mitochondrial homeostasis and regulates innate immune responses to Mycobacterium tuberculosis. Elife 9, (2020).

50. World Health Organization, Consensus meeting report: development of a Target Product Profile (TPP) and a framework for evaluation for a test for predicting progression from tuberculosis infection to active disease. (2017).

51. F. Darboe et al., Diagnostic performance of an optimized transcriptomic signature of risk of tuberculosis in cryopreserved peripheral blood mononuclear cells. Tuberculosis (Edinb) 108, 124–126 (2018).

52. S. T. Malherbe et al., Persisting positron emission tomography lesion activity and Mycobacterium tuberculosis mRNA after tuberculosis cure. Nat Med 22, 1094–1100 (2016).

53. V. Kaipilyawar et al., Development and validation of a parsimonious TB gene signature using the digital NanoString nCounter platform. Clin Infect Dis, (2022).

54. A. M. Cadena, S. M. Fortune, J. L. Flynn, Heterogeneity in tuberculosis. Nat Rev Immunol 17, 691–702 (2017).

55. M. Opperman et al., Chronological Metabolic Response to Intensive Phase TB Therapy in Patients with Cured and Failed Treatment Outcomes. ACS Infect Dis 7, 1859–1869 (2021).

56. W. A. Haynes et al., Empowering Multi-Cohort Gene Expression Analysis to Increase Reproducibility. Pac Symp Biocomput 22, 144–153 (2017).

57. S. Chen, Y. Zhou, Y. Chen, J. Gu, fastp: an ultra-fast all-in-one FASTQ preprocessor. Bioinformatics 34, i884–i890 (2018).

58. R. Patro, G. Duggal, M. I. Love, R. A. Irizarry, C. Kingsford, Salmon provides fast and bias-aware quantification of transcript expression. Nat Methods 14, 417–419 (2017).

59. M. D. Ritchie, E. R. Holzinger, R. Li, S. A. Pendergrass, D. Kim, Methods of integrating data to uncover genotype-phenotype interactions. Nat Rev Genet 16, 85–97 (2015).

60. B. Phipson, S. Lee, I. J. Majewski, W. S. Alexander, G. K. Smyth, Robust Hyperparameter Estimation Protects against Hypervariable Genes and Improves Power to Detect Differential Expression. Ann Appl Stat 10, 946–963 (2016).

61. M. D. Robinson, A. Oshlack, A scaling normalization method for differential expression analysis of RNA-seq data. Genome Biol 11, R25 (2010).

62. I. K. Jordan, L. Marino-Ramirez, Y. I. Wolf, E. V. Koonin, Conservation and coevolution in the scale-free human gene coexpression network. Mol Biol Evol 21, 2058–2070 (2004).

63. M. V. Kuleshov et al., Enrichr: a comprehensive gene set enrichment analysis web server 2016 update. Nucleic Acids Res 44, W90–97 (2016).

64. B. Efron, T. Hastie, I. Johnstone, R. Tibshirani, Least angle regression. The Annals of Statistics 32, (2004).

65. N. Meinshausen, P. Bühlmann, Stability selection. Journal of the Royal Statistical Society: Series B (Statistical Methodology) 72, 417–473 (2010).

66. S. Varma, R. Simon, Bias in error estimation when using cross-validation for model selection. BMC Bioinformatics 7, 91 (2006).

67. J. Bergstra, Y. Bengio, Random search for hyper-parameter optimization. Journal of machine learning research 13, (2012).

68. G. S. Ying, M. G. Maguire, R. J. Glynn, B. Rosner, Calculating Sensitivity, Specificity, and Predictive Values for Correlated Eye Data. Invest Ophthalmol Vis Sci 61, 29 (2020).

69. P. Massart, The Tight Constant in the Dvoretzky-Kiefer-Wolfowitz Inequality. The Annals of Probability 18, 1269–1283 (1990).

