## Supplemental figures and tables for "Common gene signature model discovery and systematic validation for TB prognosis and response to treatment"

### Supplementary materials

#### Supplementary Table 1

Cohort lists used for model training and validation. The first list contained TB-related datasets and used for training and validation analyses, along with accession IDs, platforms, and which disease comparisons each dataset was used in. The second list contained respiratory viral infection datasets for model validation. (The table is in a separated excel file)

#### Supplementary Table 2

**Pathway enrichment analysis of the common gene signatures.** Results from gene enrichment analysis on 45 candidate genes using REACTOME gene sets

| **Term** | **Overlap** | **P-value** | **Adjusted P-value** | **Genes** |
| --- | --- | --- | --- | --- |
| Interferon Signaling | 9/196 | 4.53E-10 | 1.14E-07 | GBP5, RSAD2, STAT1, JAK2, GBP1, FCGR1B, IFIT3, GBP4, IFIT2 |
| Immune System | 18/1547 | 2.03E-09 | 2.55E-07 | C1QB, CD274, GBP5, DUSP3, RSAD2, STAT1, LY96, MAPK14, IFIT3, IFIT2, AIM2, CD19, FBXO6, JAK2, TLR5, GBP1, FCGR1B, GBP4 |
| Interferon gamma signaling | 6/93 | 6.04E-08 | 5.08E-06 | GBP5, STAT1, JAK2, GBP1, FCGR1B, GBP4 |
| Cytokine Signaling in Immune system | 9/620 | 8.15E-06 | 5.13E-04 | GBP5, RSAD2, STAT1, JAK2, GBP1, FCGR1B, IFIT3, GBP4, IFIT2 |
| Interferon alpha/beta signaling | 4/68 | 1.64E-05 | 8.26E-04 | RSAD2, STAT1, IFIT3, IFIT2 |
| Interleukin-6 signaling | 2/11 | 2.69E-04 | 0.009873 | STAT1, JAK2 |
| Toll-Like Receptors Cascades | 4/140 | 2.74E-04 | 0.009873 | DUSP3, LY96, MAPK14, TLR5 |

#### Supplementary Table 3

5-fold nested cross-validation performance metrics among 7 selected machine-learning approaches using the pooled discovery datasets (27 cohorts, datapoints *n* = 2914). Outer CV AUROC presented graphically in Supplementary Figure 6.

| **ML type** | **Cross Validation fold** | **Inner CV mean squared error** | **Outer CV mean squared error** | **Outer CV r2** | **Outer CV AUROC** | **# features** |
| --- | --- | --- | --- | --- | --- | --- |
| Support vector machine | Fold 1 | 0.12535 | 0.134806736 | 0.422078385 | 0.888469 | 35 |
|  | Fold 2 | 0.12223 | 0.16202878 | 0.271557799 | 0.835513 | 37 |
|  | Fold 3 | 0.1232 | 0.145485714 | 0.368675246 | 0.882899 | 36 |
|  | Fold 4 | 0.12161 | 0.121276732 | 0.449320213 | 0.896788 | 42 |
|  | Fold 5 | 0.12477 | 0.139605354 | 0.353392267 | 0.87495 | 33 |
| Random forest | Fold 1 | 0.10824 | 0.115712122 | 0.47189422 | 0.900731 | 35 |
|  | Fold 2 | 0.11112 | 0.104998793 | 0.531890915 | 0.923051 | 37 |
|  | Fold 3 | 0.10725 | 0.102516937 | 0.529520189 | 0.921499 | 36 |
|  | Fold 4 | 0.10975 | 0.105128641 | 0.519461024 | 0.920225 | 42 |
|  | Fold 5 | 0.11133 | 0.107287322 | 0.505775306 | 0.914664 | 33 |
| Elastic net | Fold 1 | 0.16553 | 0.177626051 | 0.177432507 | 0.811288 | 35 |
|  | Fold 2 | 0.16847 | 0.167222478 | 0.223634104 | 0.847927 | 37 |
|  | Fold 3 | 0.16646 | 0.173145045 | 0.197150056 | 0.826173 | 36 |
|  | Fold 4 | 0.16898 | 0.171061369 | 0.20658821 | 0.839441 | 42 |
|  | Fold 5 | 0.16886 | 0.162171398 | 0.246372257 | 0.832584 | 33 |
| Adaptive boosting | Fold 1 | 0.13479 | 0.146447768 | 0.331186836 | 0.8472 | 35 |
|  | Fold 2 | 0.14148 | 0.136208534 | 0.389743832 | 0.872539 | 37 |
|  | Fold 3 | 0.1333 | 0.132927767 | 0.395496792 | 0.876505 | 36 |
|  | Fold 4 | 0.13346 | 0.141352288 | 0.346929548 | 0.861884 | 42 |
|  | Fold 5 | 0.13776 | 0.135681254 | 0.385407643 | 0.870585 | 33 |
| Partial least squares | Fold 1 | 0.16383 | 0.178489261 | 0.174899137 | 0.809612 | 35 |
|  | Fold 2 | 0.1677 | 0.166340333 | 0.228971917 | 0.851073 | 37 |
|  | Fold 3 | 0.1656 | 0.173703028 | 0.196260158 | 0.821547 | 36 |
|  | Fold 4 | 0.16757 | 0.169972661 | 0.21199362 | 0.836877 | 42 |
|  | Fold 5 | 0.16837 | 0.163176985 | 0.241745793 | 0.822385 | 33 |
| Multilayer perceptron | Fold 1 | 0.11068 | 0.120884813 | 0.442106526 | 0.895058 | 35 |
|  | Fold 2 | 0.11319 | 0.113109607 | 0.492675002 | 0.881656 | 37 |
|  | Fold 3 | 0.11321 | 0.112396706 | 0.484707383 | 0.896247 | 36 |
|  | Fold 4 | 0.11772 | 0.10990995 | 0.49043002 | 0.903789 | 42 |
|  | Fold 5 | 0.11553 | 0.116177792 | 0.485655834 | 0.907721 | 33 |
| Extreme gradient boosting | Fold 1 | 0.10796 | 0.141314945 | 0.349845073 | 0.854219 | 35 |
|  | Fold 2 | 0.11344 | 0.119385948 | 0.455040281 | 0.899911 | 37 |
|  | Fold 3 | 0.11076 | 0.140167115 | 0.389811924 | 0.874422 | 36 |
|  | Fold 4 | 0.11200 | 0.122377948 | 0.444486009 | 0.893244 | 42 |
|  | Fold 5 | 0.11381 | 0.132638969 | 0.423787823 | 0.891314 | 33 |

#### Supplementary Table 4

The parameters of the final models trained by the pooled discovery dataset (27 cohorts)

| **Model** | **parameters** | **# features** | **Features (paired genes)** | **# genes selected in features** | **Genes** |
| --- | --- | --- | --- | --- | --- |
| Full model | RandomForestRegressor (bootstrap=False, max_depth=30, max_features='sqrt', n_estimators=700, random_state=1) | 41 | SPOCK2_DUSP3, SPOCK2_STAT1, CD19_SERPING1, CD19_LAP3, FBXO6_C1QB, C1QB_SLC6A12, C1QB_RTP4, CD19_DUSP3, GK_LRRK2, SPOCK2_IFIT2, PSTPIP2_SERPING1, ZNF438_LHFPL2, SPOCK2_CD5, FBXO6_FCGR1B, CD274_MAPK14, AIM2_SERPING1, BATF2_ANKRD22, LRRK2_LMNB1, CD274_APOL6, HP_C1QB, PSTPIP2_LHFPL2, SAMD9L_LAP3, CD274_JAK2, NELL2_CD5, BATF2_C1QB, CASP5_FCGR1B, GBP5_GBP4, CD274_ZNF438, CD274_P2RY14, TIMM10_FBXO6, DUSP3_LY96, FBXO6_GBP1, TLR5_ZNF438, ADM_IFIT2, RSAD2_IFIT3, RSAD2_GBP5, RSAD2_FCGR1B, LRRK2_KCNJ15, GBP4_LAP3, FBXO6_VAMP5, ZNF438_KCNJ15 | 42 | ADM, AIM2, ANKRD22, APOL6, BATF2, C1QB, CASP5, CD19, CD274, CD5, DUSP3, GBP1, GBP4, GBP5, FBXO6, FCGR1B, GK, HP, IFIT2, IFIT3, JAK2, KCNJ15, LAP3, LHFPL2, LMNB1, LRRK2, LY96, MAPK14, NELL2, P2RY14, PSTPIP2, RSAD2, RTP4, SAMD9L, SERPING1, SLC6A12, SPOCK2, STAT1, TIMM10, TLR5, VAMP5, ZNF438 |
| Reduced model | RandomForestRegressor (bootstrap=False, max_depth=90, max_features='sqrt', n_estimators=1400, random_state=1) | 12 | FBXO6_VAMP5, LRRK2_LMNB1, BATF2_ANKRD22, SPOCK2_DUSP3, CD274_NELL2, GBP5_GBP4, IFIT2_ADM, ZNF438_FCGR1B, NELL2_CD5, CD274_APOL6, SPOCK2_CD5, IFIT2_SPOCK2 | 18 | ADM, ANKRD22, APOL6, BATF2, CD274, CD5, DUSP3, GBP4, GBP5, FBXO6, FCGR1B, LMNB1, LRRK2, IFIT2, NELL2, SPOCK2, VAMP5, ZNF438 |

#### Supplementary Table 5

Prognostic performance of the new models developed in this report and published previously for incipient TB, stratified by time interval to disease, using cut-offs identified by the maximal Youden Index based on the best tradeoff between sensitivity and specificity from each ROC. Positive and negative predictive values (PPVs/NPVs) were calculated when assuming 2% pre-test probability. The performance metrics are presented with 95 % confidence interval.

|  | Time interval | Sensitivity | Specificity | PPV | NPV |
| --- | --- | --- | --- | --- | --- |
| Full model | < 3m to disease | 0.873 (0.815 - 0.931) | 0.917 (0.899 - 0.935) | 0.177 (0.145 - 0.209) | 0.997 (0.996 - 0.998) |
|  | < 6m to disease | 0.821 (0.764 - 0.878) | 0.838 (0.813 - 0.862) | 0.093 (0.081 - 0.106) | 0.996 (0.995 - 0.997) |
|  | < 12m to disease | 0.786 (0.734 - 0.839) | 0.803 (0.777 - 0.830) | 0.075 (0.066 - 0.085) | 0.995 (0.994 - 0.995) |
|  | < 18m to disease | 0.773 (0.723 - 0.823) | 0.783 (0.756 - 0.810) | 0.068 (0.060 - 0.076) | 0.994 (0.993 - 0.995) |
|  | < 24m to disease | 0.747 (0.697 - 0.796) | 0.783 (0.756 - 0.810) | 0.066 (0.058 - 0.073) | 0.993 (0.993 - 0.994) |
|  | < 30m to disease | 0.742 (0.693 - 0.792) | 0.783 (0.756 - 0.810) | 0.065 (0.058 - 0.073) | 0.993 (0.993 - 0.994) |
| Reduced model | < 3m to disease | 0.833 (0.768 - 0.898) | 0.916 (0.898 - 0.934) | 0.168 (0.138 - 0.199) | 0.996 (0.995 - 0.997) |
|  | < 6m to disease | 0.740 (0.675 - 0.805) | 0.869 (0.847 - 0.892) | 0.104 (0.088 - 0.119) | 0.994 (0.993 - 0.995) |
|  | < 12m to disease | 0.748 (0.692 - 0.804) | 0.810 (0.784 - 0.836) | 0.074 (0.065 - 0.084) | 0.994 (0.993 - 0.995) |
|  | < 18m to disease | 0.784 (0.735 - 0.834) | 0.749 (0.720 - 0.778) | 0.060 (0.053 - 0.066) | 0.994 (0.993 - 0.995) |
|  | < 24m to disease | 0.771 (0.722 - 0.819) | 0.749 (0.720 - 0.778) | 0.059 (0.053 - 0.065) | 0.994 (0.993 - 0.995) |
|  | < 30m to disease | 0.773 (0.725 - 0.820) | 0.749 (0.720 - 0.778) | 0.059 (0.053 - 0.065) | 0.994 (0.993 - 0.995) |
| Sweeney 3 | < 3m to disease | 0.714 (0.635 - 0.793) | 0.464 (0.431 - 0.497) | 0.026 (0.025 - 0.028) | 0.988 (0.986 - 0.989) |
|  | < 6m to disease | 0.751 (0.687 - 0.816) | 0.401 (0.369 - 0.434) | 0.025 (0.023 - 0.026) | 0.988 (0.986 - 0.989) |
|  | < 12m to disease | 0.752 (0.697 - 0.807) | 0.401 (0.369 - 0.434) | 0.025 (0.024 - 0.026) | 0.988 (0.986 - 0.989) |
|  | < 18m to disease | 0.725 (0.672 - 0.778) | 0.401 (0.369 - 0.434) | 0.024 (0.023 - 0.026) | 0.986 (0.985 - 0.987) |
|  | < 24m to disease | 0.705 (0.653 - 0.758) | 0.401 (0.369 - 0.434) | 0.023 (0.022 - 0.025) | 0.985 (0.984 - 0.986) |
|  | < 30m to disease | 0.709 (0.658 - 0.761) | 0.401 (0.369 - 0.434) | 0.024 (0.022 - 0.025) | 0.985 (0.984 - 0.987) |
| RISK 6 | < 3m to disease | 0.857 (0.796 - 0.918) | 0.865 (0.842 - 0.887) | 0.115 (0.098 - 0.131) | 0.997 (0.996 - 0.998) |
|  | < 6m to disease | 0.798 (0.738 - 0.858) | 0.836 (0.812 - 0.861) | 0.090 (0.078 - 0.103) | 0.995 (0.994 - 0.996) |
|  | < 12m to disease | 0.752 (0.697 - 0.807) | 0.836 (0.812 - 0.861) | 0.086 (0.074 - 0.097) | 0.994 (0.993 - 0.995) |
|  | < 18m to disease | 0.714 (0.660 - 0.768) | 0.836 (0.812 - 0.861) | 0.082 (0.071 - 0.093) | 0.993 (0.992 - 0.994) |
|  | < 24m to disease | 0.740 (0.689 - 0.790) | 0.761 (0.733 - 0.790) | 0.059 (0.053 - 0.066) | 0.993 (0.992 - 0.994) |
|  | < 30m to disease | 0.732 (0.682 - 0.783) | 0.761 (0.733 - 0.790) | 0.059 (0.052 - 0.065) | 0.993 (0.992 - 0.994) |
| BATF2 | < 3m to disease | 0.738 (0.661 - 0.815) | 0.594 (0.562 - 0.627) | 0.036 (0.033 - 0.039) | 0.991 (0.990 - 0.992) |
|  | < 6m to disease | 0.717 (0.650 - 0.784) | 0.555 (0.522 - 0.587) | 0.032 (0.029 - 0.034) | 0.990 (0.989 - 0.991) |
|  | < 12m to disease | 0.662 (0.602 - 0.723) | 0.549 (0.516 - 0.582) | 0.029 (0.027 - 0.031) | 0.988 (0.987 - 0.989) |
|  | < 18m to disease | 0.721 (0.668 - 0.775) | 0.476 (0.443 - 0.509) | 0.027 (0.026 - 0.029) | 0.988 (0.987 - 0.989) |
|  | < 24m to disease | 0.709 (0.657 - 0.761) | 0.472 (0.439 - 0.505) | 0.027 (0.025 - 0.028) | 0.988 (0.987 - 0.989) |
|  | < 30m to disease | 0.702 (0.651 - 0.754) | 0.472 (0.439 - 0.505) | 0.026 (0.025 - 0.028) | 0.987 (0.986 - 0.988) |
| Suliman 4 | < 3m to disease | 0.841 (0.777 - 0.905) | 0.831 (0.806 - 0.855) | 0.092 (0.080 - 0.104) | 0.996 (0.995 - 0.997) |
|  | < 6m to disease | 0.746 (0.681 - 0.811) | 0.831 (0.806 - 0.855) | 0.082 (0.071 - 0.094) | 0.994 (0.993 - 0.995) |
|  | < 12m to disease | 0.667 (0.606 - 0.727) | 0.831 (0.806 - 0.855) | 0.074 (0.064 - 0.084) | 0.992 (0.991 - 0.993) |
|  | < 18m to disease | 0.617 (0.559 - 0.675) | 0.831 (0.806 - 0.855) | 0.069 (0.060 - 0.079) | 0.991 (0.990 - 0.992) |
|  | < 24m to disease | 0.716 (0.664 - 0.767) | 0.670 (0.639 - 0.702) | 0.042 (0.039 - 0.046) | 0.991 (0.991 - 0.992) |
|  | < 30m to disease | 0.712 (0.661 - 0.764) | 0.670 (0.639 - 0.702) | 0.042 (0.038 - 0.046) | 0.991 (0.991 - 0.992) |

#### Supplementary Table 6

Prognostic performance of the models developed in this report and published previously for incipient TB, stratified by time interval to disease, using the predetermined cutoffs defined by 2 standard deviations (SDs) above the mean of the control group (latent TB infection without progression during follow-up). Positive and negative predictive values (PPVs/NPVs) were calculated when assuming 2% pre-test probability. The performance metrics presented with 95 % confidence interval.

|  | Time interval | Sensitivity | Specificity | PPV | NPV |
| --- | --- | --- | --- | --- | --- |
| Full model | < 3m to disease | 0.667 (0.584 - 0.749) | 0.965 (0.953 - 0.977) | 0.279 (0.209 - 0.348) | 0.993 (0.991 - 0.995) |
|  | < 6m to disease | 0.561 (0.487 - 0.635) | 0.965 (0.953 - 0.977) | 0.245 (0.181 - 0.309) | 0.991 (0.989 - 0.992) |
|  | < 12m to disease | 0.487 (0.423 - 0.551) | 0.965 (0.953 - 0.977) | 0.220 (0.161 - 0.279) | 0.989 (0.988 - 0.990) |
|  | < 18m to disease | 0.446 (0.387 - 0.506) | 0.965 (0.953 - 0.977) | 0.205 (0.149 - 0.262) | 0.988 (0.987 - 0.990) |
|  | < 24m to disease | 0.418 (0.361 - 0.474) | 0.965 (0.953 - 0.977) | 0.195 (0.141 - 0.249) | 0.988 (0.987 - 0.989) |
|  | < 30m to disease | 0.411 (0.356 - 0.467) | 0.965 (0.953 - 0.977) | 0.192 (0.139 - 0.246) | 0.988 (0.987 - 0.989) |
| Reduced model | < 3m to disease | 0.690 (0.610 - 0.771) | 0.967 (0.955 - 0.979) | 0.300 (0.224 - 0.375) | 0.994 (0.992 - 0.995) |
|  | < 6m to disease | 0.566 (0.493 - 0.640) | 0.967 (0.955 - 0.979) | 0.260 (0.191 - 0.329) | 0.991 (0.990 - 0.992) |
|  | < 12m to disease | 0.483 (0.419 - 0.547) | 0.967 (0.955 - 0.979) | 0.230 (0.167 - 0.294) | 0.989 (0.988 - 0.990) |
|  | < 18m to disease | 0.439 (0.379 - 0.498) | 0.967 (0.955 - 0.979) | 0.214 (0.153 - 0.274) | 0.988 (0.987 - 0.989) |
|  | < 24m to disease | 0.418 (0.361 - 0.474) | 0.967 (0.955 - 0.979) | 0.206 (0.147 - 0.264) | 0.988 (0.987 - 0.989) |
|  | < 30m to disease | 0.411 (0.356 - 0.467) | 0.967 (0.955 - 0.979) | 0.203 (0.145 - 0.261) | 0.988 (0.987 - 0.989) |
| Sweeney 3 | < 3m to disease | 0.048 (0.010 - 0.085) | 0.990 (0.983 - 0.996) | 0.087 (0.035 - 0.138) | 0.981 (0.980 - 0.981) |
|  | < 6m to disease | 0.046 (0.015 - 0.078) | 0.990 (0.983 - 0.996) | 0.084 (0.034 - 0.135) | 0.981 (0.980 - 0.981) |
|  | < 12m to disease | 0.056 (0.026 - 0.085) | 0.990 (0.983 - 0.996) | 0.100 (0.041 - 0.158) | 0.981 (0.980 - 0.981) |
|  | < 18m to disease | 0.048 (0.023 - 0.074) | 0.990 (0.983 - 0.996) | 0.088 (0.036 - 0.140) | 0.981 (0.980 - 0.981) |
|  | < 24m to disease | 0.045 (0.021 - 0.068) | 0.990 (0.983 - 0.996) | 0.082 (0.033 - 0.130) | 0.981 (0.980 - 0.981) |
|  | < 30m to disease | 0.043 (0.020 - 0.067) | 0.990 (0.983 - 0.996) | 0.080 (0.032 - 0.128) | 0.981 (0.980 - 0.981) |
| RISK 6 | < 3m to disease | 0.643 (0.559 - 0.727) | 0.965 (0.953 - 0.977) | 0.271 (0.203 - 0.340) | 0.993 (0.991 - 0.994) |
|  | < 6m to disease | 0.549 (0.475 - 0.623) | 0.965 (0.953 - 0.977) | 0.241 (0.178 - 0.305) | 0.991 (0.989 - 0.992) |
|  | < 12m to disease | 0.474 (0.410 - 0.538) | 0.965 (0.953 - 0.977) | 0.216 (0.157 - 0.274) | 0.989 (0.988 - 0.990) |
|  | < 18m to disease | 0.428 (0.368 - 0.487) | 0.965 (0.953 - 0.977) | 0.199 (0.143 - 0.254) | 0.988 (0.987 - 0.989) |
|  | < 24m to disease | 0.408 (0.351 - 0.464) | 0.965 (0.953 - 0.977) | 0.191 (0.138 - 0.244) | 0.988 (0.987 - 0.989) |
|  | < 30m to disease | 0.401 (0.346 - 0.457) | 0.965 (0.953 - 0.977) | 0.189 (0.136 - 0.242) | 0.987 (0.986 - 0.989) |
| BATF2 | < 3m to disease | 0.016 (0.000 - 0.038) | 0.967 (0.955 - 0.979) | 0.010 (0.006 - 0.013) | 0.980 (0.979 - 0.980) |
|  | < 6m to disease | 0.023 (0.001 - 0.046) | 0.967 (0.955 - 0.979) | 0.014 (0.009 - 0.019) | 0.980 (0.979 - 0.980) |
|  | < 12m to disease | 0.030 (0.008 - 0.052) | 0.967 (0.955 - 0.979) | 0.018 (0.012 - 0.025) | 0.980 (0.979 - 0.980) |
|  | < 18m to disease | 0.026 (0.007 - 0.045) | 0.967 (0.955 - 0.979) | 0.016 (0.010 - 0.021) | 0.980 (0.979 - 0.980) |
|  | < 24m to disease | 0.024 (0.006 - 0.042) | 0.967 (0.955 - 0.979) | 0.015 (0.009 - 0.020) | 0.980 (0.979 - 0.980) |
|  | < 30m to disease | 0.023 (0.006 - 0.041) | 0.967 (0.955 - 0.979) | 0.014 (0.009 - 0.019) | 0.980 (0.979 - 0.980) |
| Suliman 4 | < 3m to disease | 0.563 (0.477 - 0.650) | 0.966 (0.954 - 0.978) | 0.252 (0.186 - 0.319) | 0.991 (0.989 - 0.993) |
|  | < 6m to disease | 0.491 (0.417 - 0.566) | 0.966 (0.954 - 0.978) | 0.227 (0.166 - 0.289) | 0.989 (0.988 - 0.991) |
|  | < 12m to disease | 0.419 (0.356 - 0.482) | 0.966 (0.954 - 0.978) | 0.200 (0.144 - 0.257) | 0.988 (0.987 - 0.989) |
|  | < 18m to disease | 0.379 (0.321 - 0.437) | 0.966 (0.954 - 0.978) | 0.185 (0.132 - 0.238) | 0.987 (0.986 - 0.988) |
|  | < 24m to disease | 0.360 (0.305 - 0.415) | 0.966 (0.954 - 0.978) | 0.177 (0.126 - 0.228) | 0.987 (0.986 - 0.988) |
|  | < 30m to disease | 0.355 (0.300 - 0.409) | 0.966 (0.954 - 0.978) | 0.175 (0.124 - 0.226) | 0.987 (0.986 - 0.988) |

#### Supplementary Figure 1

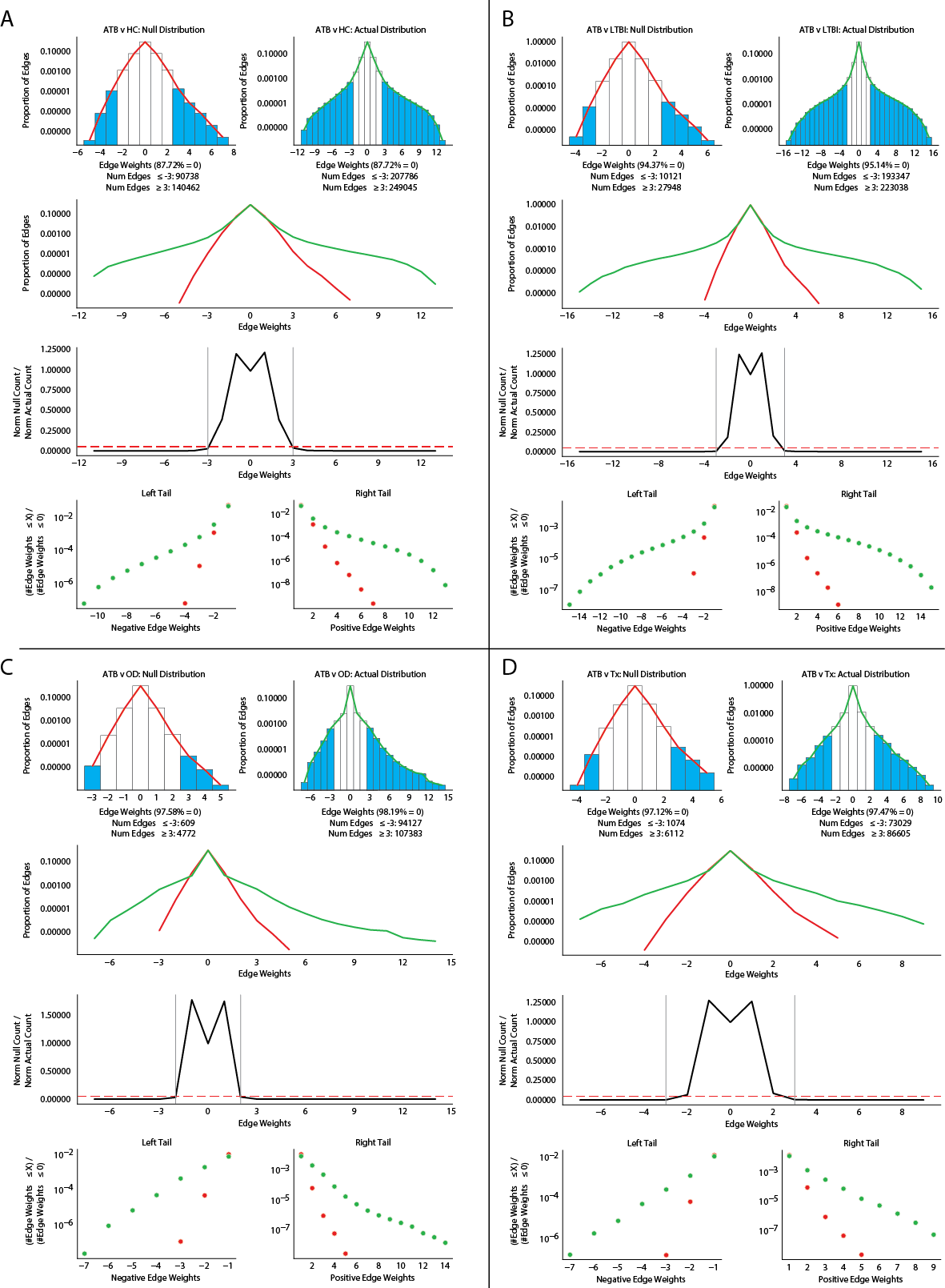
Network building parameter selection. (A-D) Null distribution of edge weights from permuting log2(Fold Change) values across cohorts and calculating dot products compared against the actual distribution of edge weights from the matrix of log2(Fold Change) values for a given disease comparison (Methods). Edge weights between -2 and 2 are likely to arise by chance and are discarded during network construction.

#### Supplementary Figure 2

Network degree distribution. Distribution of degree, weighted degree, and eigenvector centrality for the four different networks constructed from (A) ATB vs. HC, (B) ATB vs. LTBI, (C) ATB vs. OD, and (D) ATB vs. Tx disease comparisons.

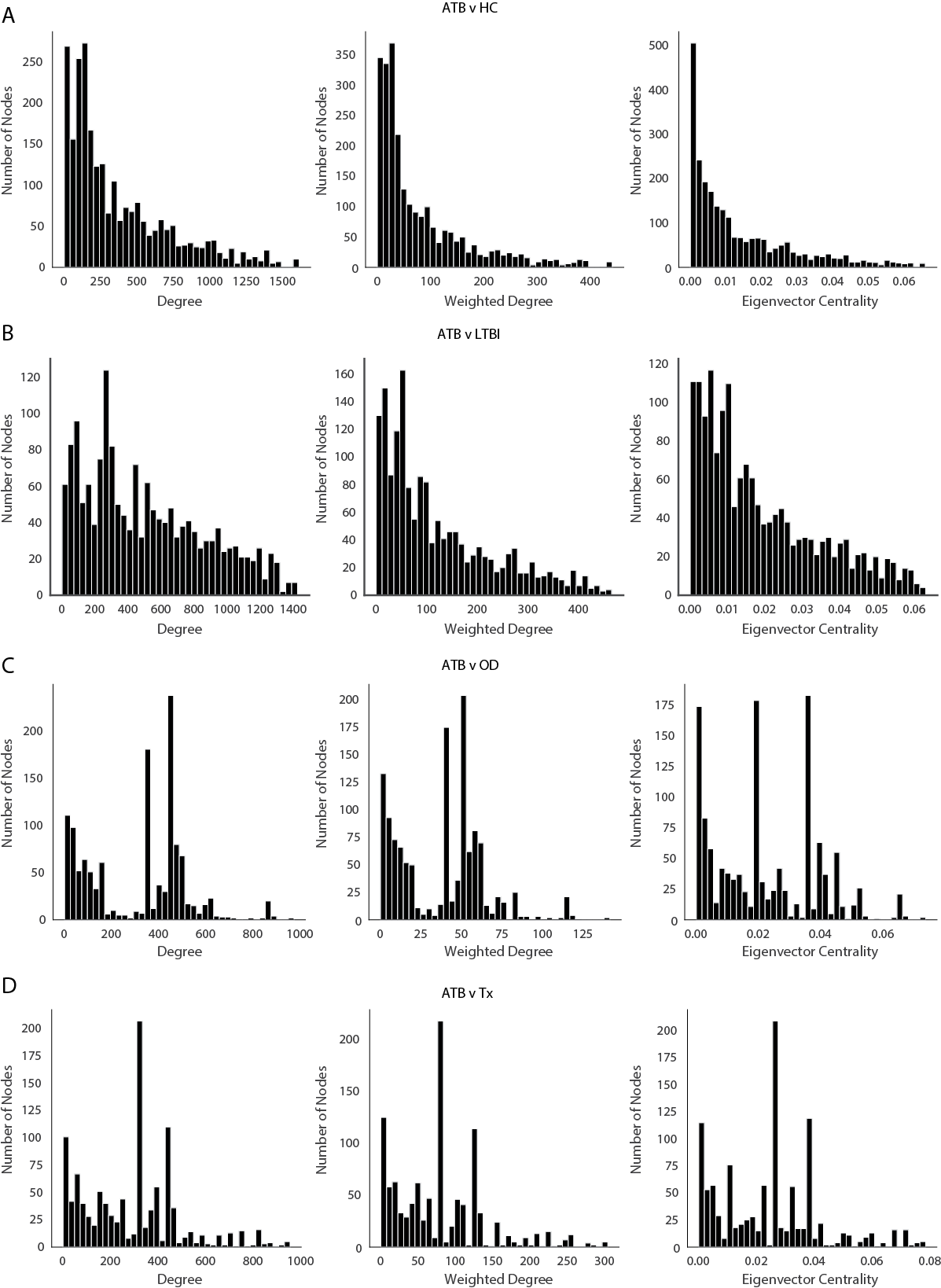

#### Supplementary Figure 3

Network degree distribution fitting. (A-D) The distribution of weighted degrees for each network was fit with a probability density function to help determine top-ranked genes to retain for the candidate gene set (Methods). (E) Overlap of top 5% of genes by weighted degree in each network.

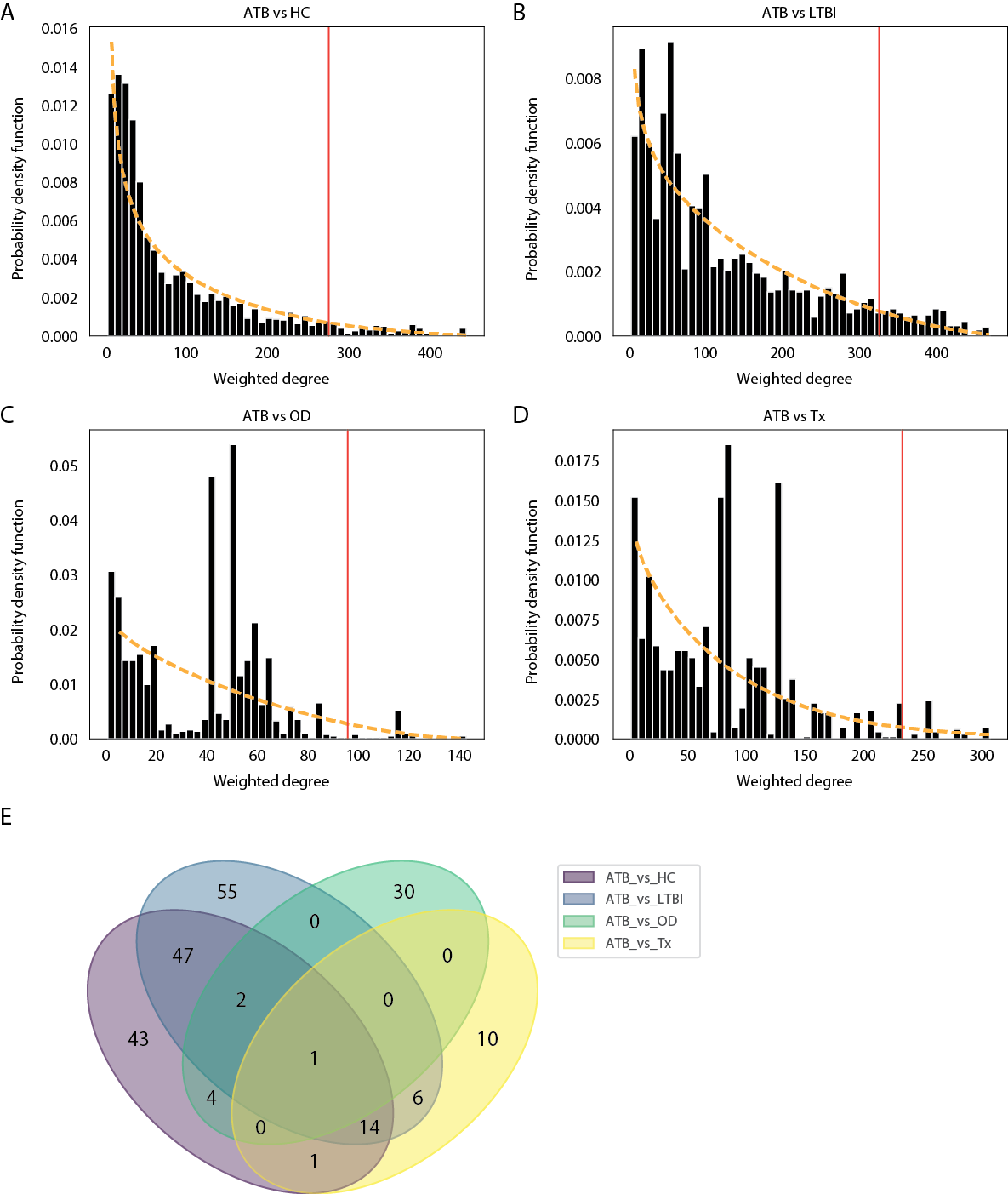

#### Supplementary Figure 4

Gene signature mapped on protein-protein interaction network. Protein-protein association network of 45 candidate genes constructed with STRING. Width of edge corresponds to edge confidence. Associations correspond to physical interactions or represent proteins that act functionally together.

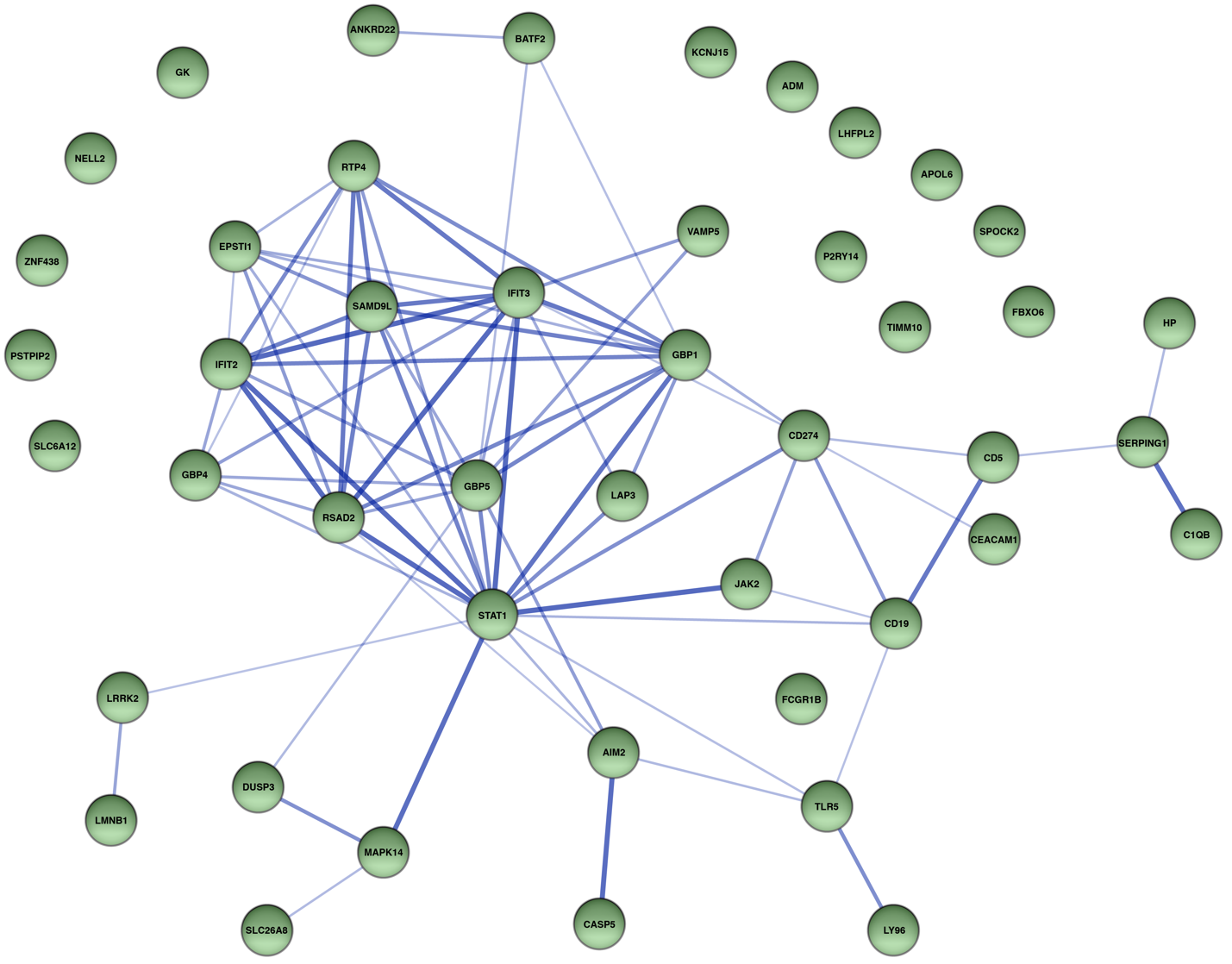

#### Supplementary Figure 5

Heatmap of gene signatures. Genes included in our 45-candidate gene set (the first column in both panels) as compared to genes in gene sets from 30 previously published gene signatures (*1, 2*). We exclude the genes (563 out of total 721 genes) only detected in one gene signature and display 155 genes present in at least two gene signatures and 3 additional genes present only in our 45-candidate gene set. The left panel displays genes detected in our candidate gene set while the right panel displays genes detected in at least two gene signatures. The numbers in parentheses indicate the number of studies identifying the gene.

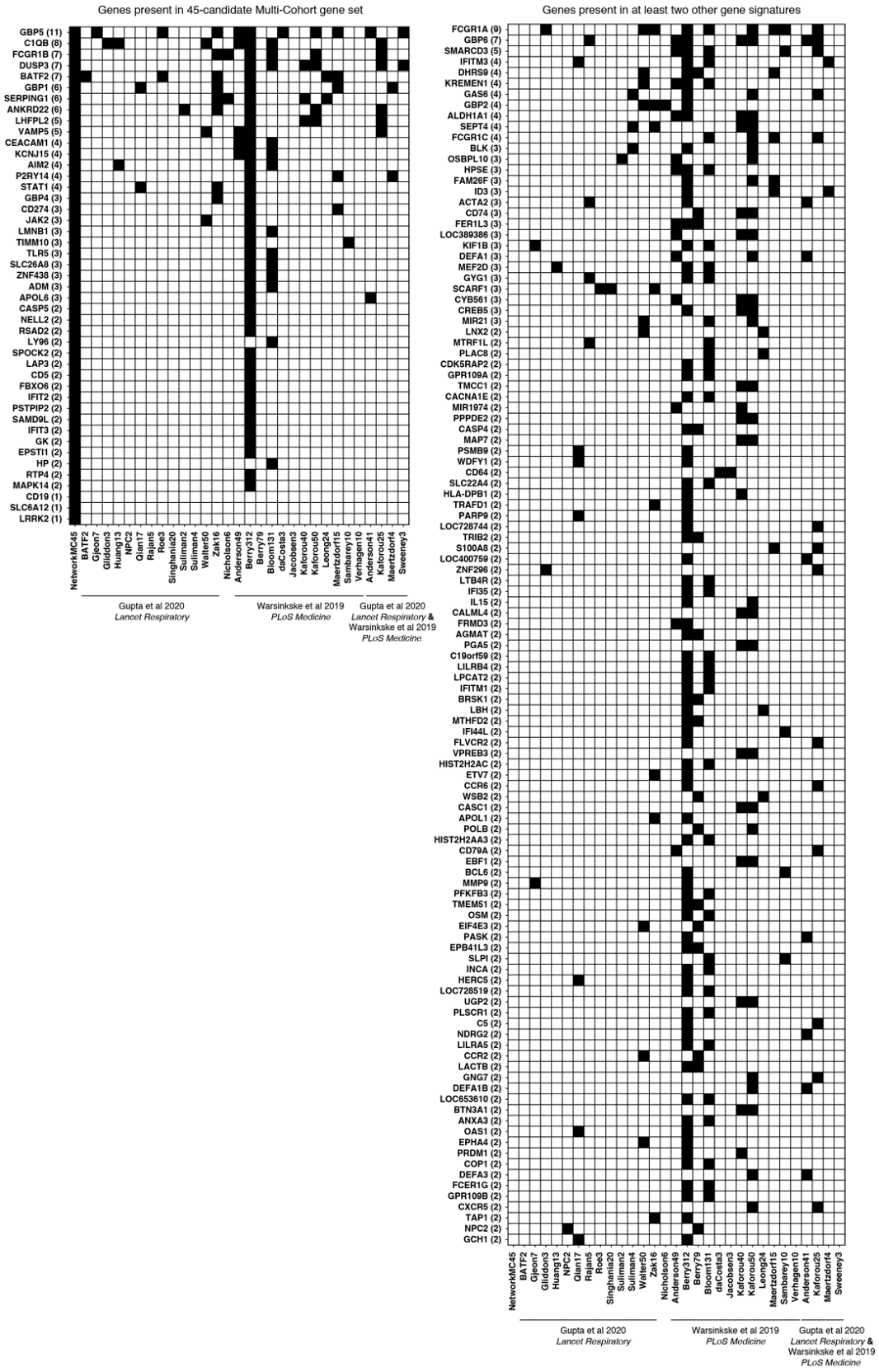

#### Supplementary Figure 6

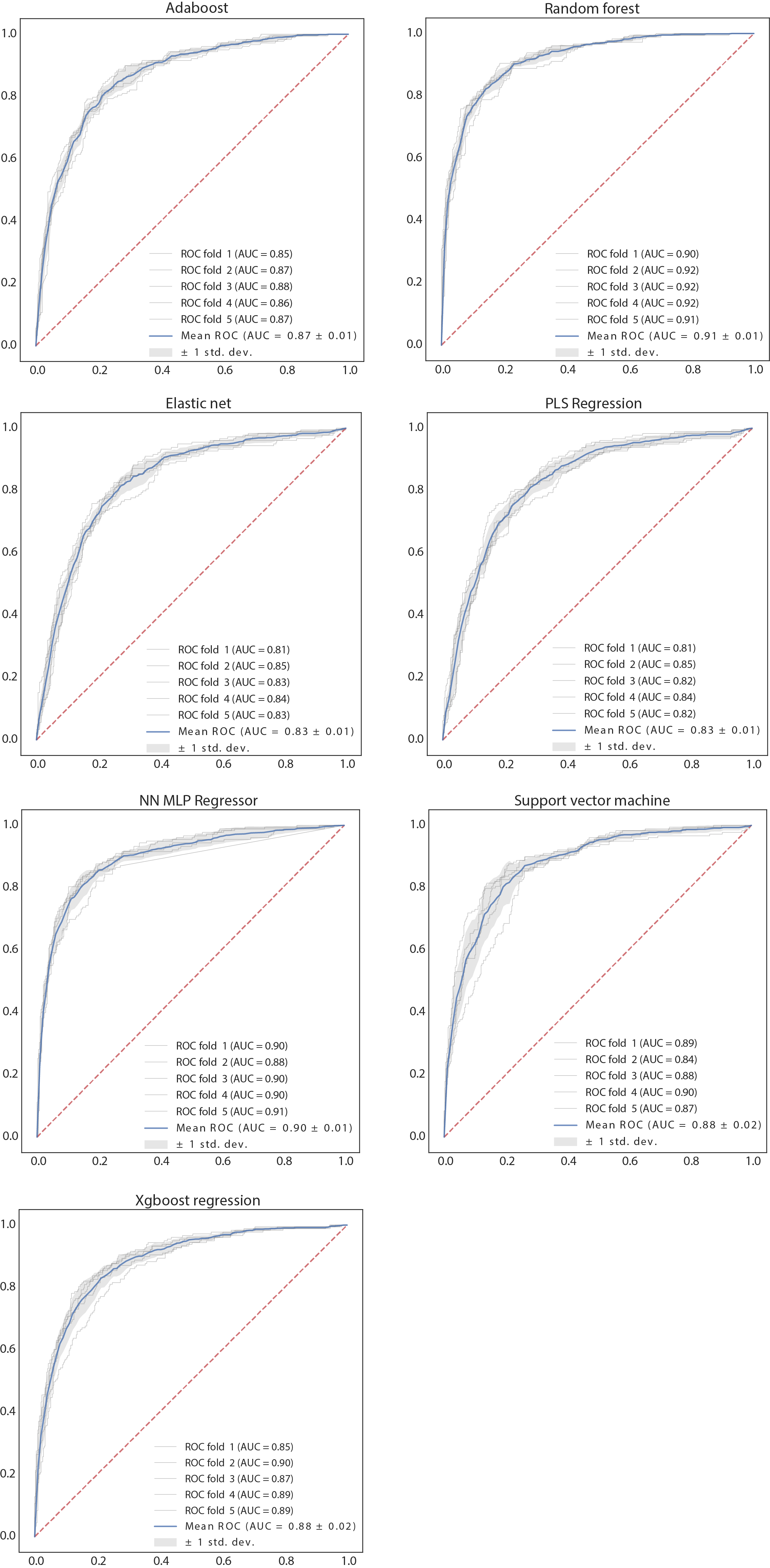
Head-to-head comparison of model performance generated from 5-fold nested cross-validation among 7 selected supervised learning models using the pooled discovery datasets (27 cohorts, datapoints *n* = 2914).

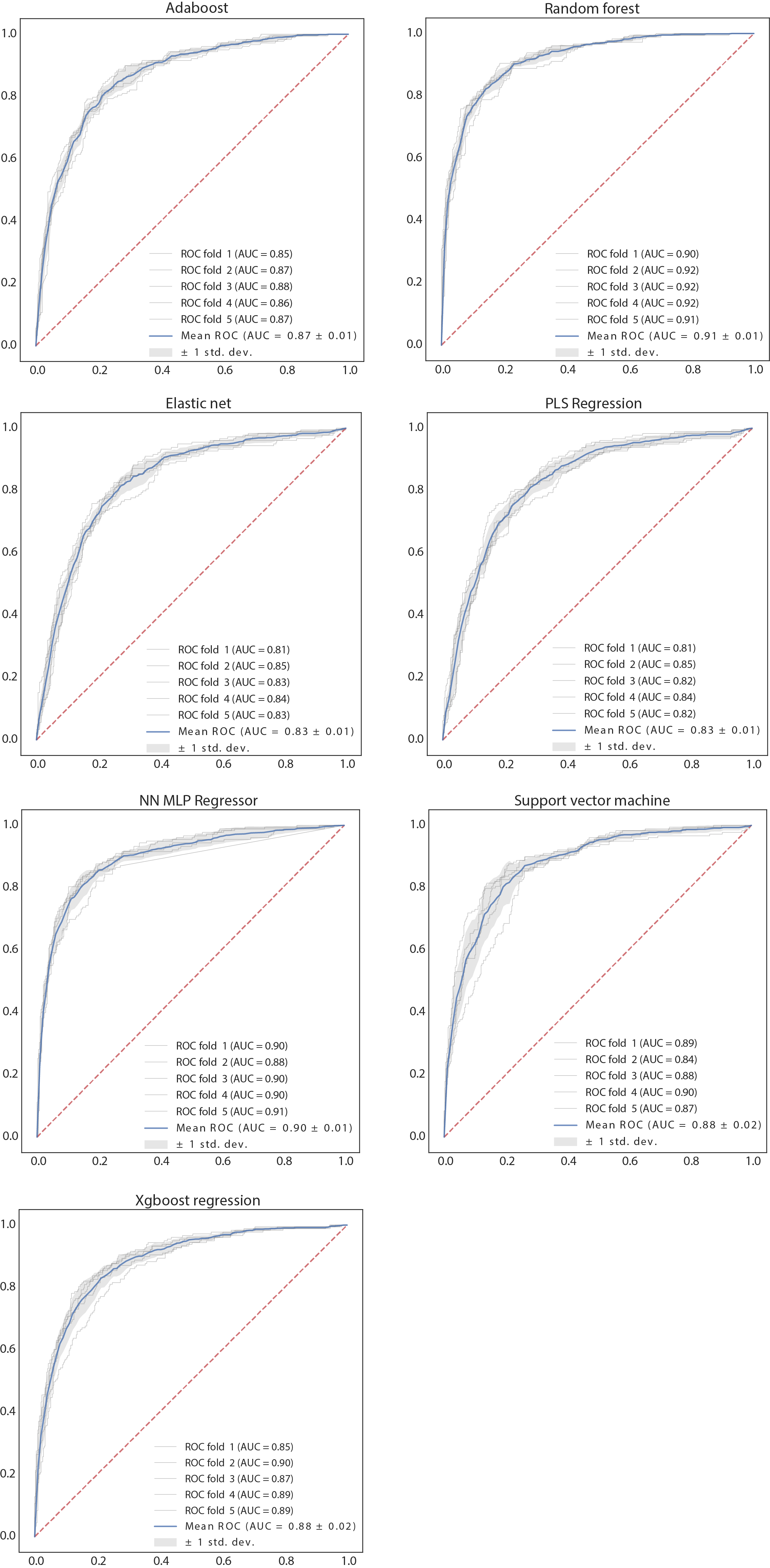
**Supplementary Figure 6**. *Continued*

#### Supplementary Figure 7

Feature down-selection by a stability analysis. (A) The stability path for each feature shown as blue line indicates the probability of a feature being selected from randomly resampling the data as a function of the regularization parameter (***λ***). Features were ranked by the maximum probability and top 20 features were listed in (B). The top 12 features were selected in our reduced model to minimize the number of features while maintaining CV performance (**Supplementary Fig. 8**).

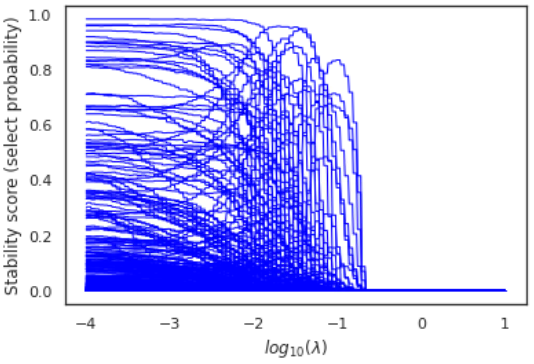
**A	B**

| **Feature (paired genes) ranking** | **Probability** |
| --- | --- |
| VAMP5_FBXO6 | 0.983 |
| LMNB1_LRRK2 | 0.965 |
| BATF2_ANKRD22 | 0.962 |
| SPOCK2_DUSP3 | 0.957 |
| CD274_NELL2 | 0.948 |
| GBP5_GBP4 | 0.941 |
| IFIT2_ADM | 0.919 |
| ZNF438_FCGR1B | 0.912 |
| NELL2_CD5 | 0.902 |
| CD274_APOL6 | 0.9 |
| SPOCK2_CD5 | 0.897 |
| IFIT2_SPOCK2 | 0.893 |
| ZNF438_KCNJ15 | 0.885 |
| GK_LRRK2 | 0.884 |
| CASP5_FCGR1B | 0.869 |
| ZNF438_CD274 | 0.86 |
| SPOCK2_STAT1 | 0.844 |
| CD19_C1QB | 0.84 |
| NELL2_C1QB | 0.835 |
| LY96_DUSP3 | 0.834 |

#### Supplementary Figure 8

Feature down-selection of the model based on AUROC with 95 % confidence intervals from 5-fold CV. The features were ranked by the stability of features selected from randomly resampling the data (**Supplementary Fig. 7**). A forward stepwise selection was used to evaluate the model CV performance by adding one feature at a time stating from the top feature. The final feature set was determined when the breakpoint of the AUROC is reached.

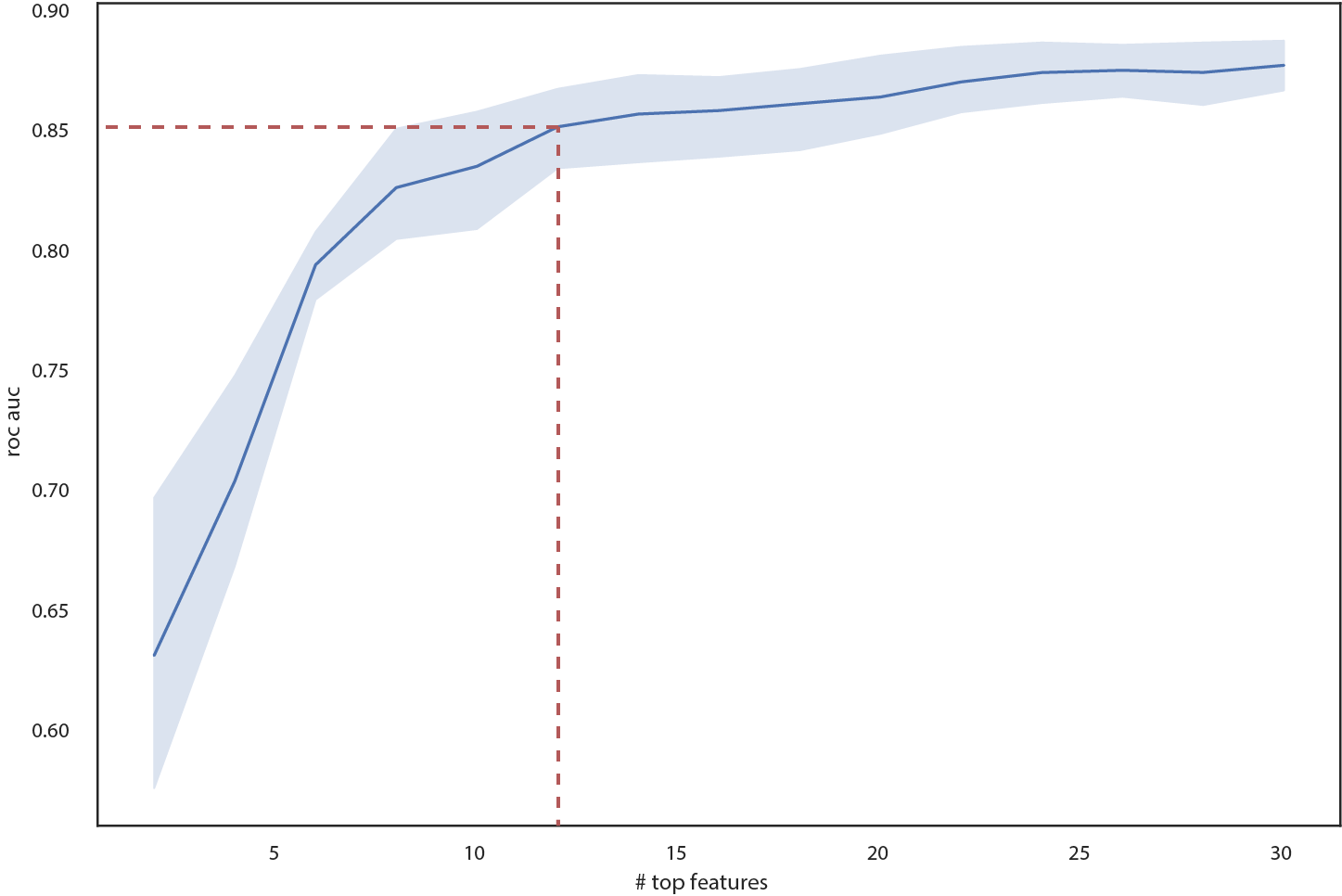

#### Supplementary Figure 9

Prognostic performance of the reduced model for incipient TB using the pooled longitudinal validation datasets. The distributions of TB scores generated by the reduced mode, stratified by categorical interval to disease, are shown in a violin plot (datapoints *n* = 1281) (left panel). ROC curves depict prognostic performance for incipient TB, stratified by time intervals to disease (< 3, < 6, <12, <18, < 24, < 30 months) (middle panel) and mutually exclusive time intervals to disease (0-3, 3-6, 6-12, 12-18, 18-24, 24-30 months) (right panel). AUC and 95% confidence intervals for each interval to disease are shown.

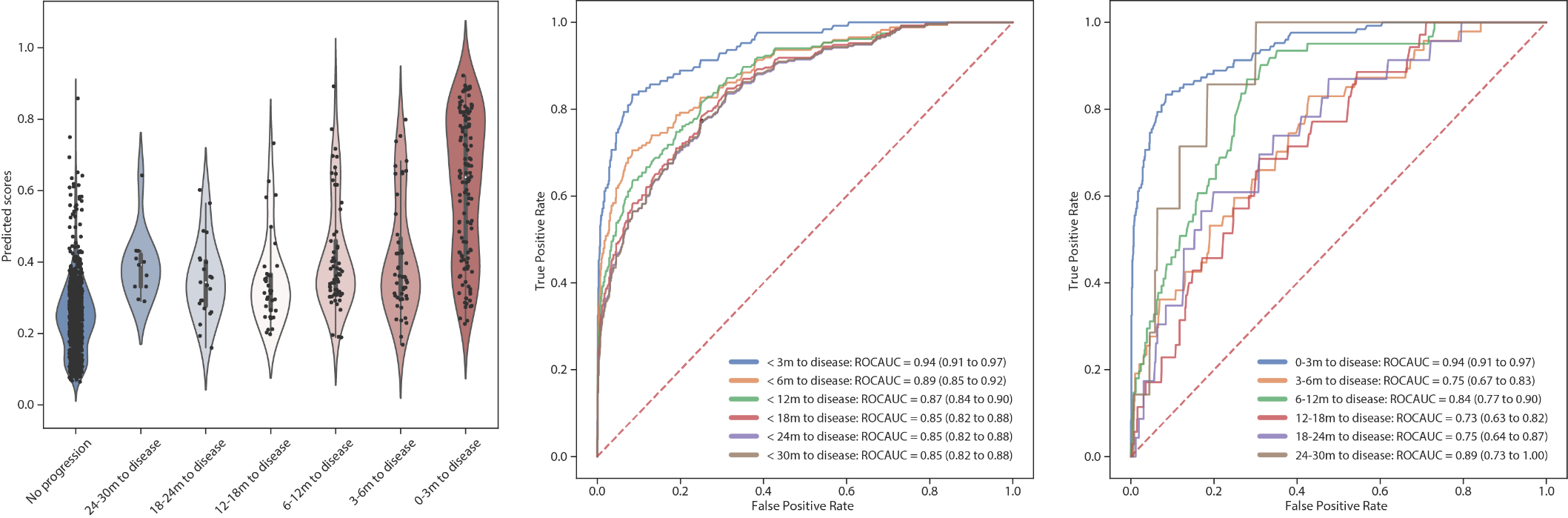

#### Supplementary Figure 10

Prognostic performance of the 4 published models(*3-6*) (A-D) for incipient TB using the pooled longitudinal validation dataset (6 TB progression studies). The distributions of TB scores, stratified by categorical interval to disease, are shown in a violin plot (datapoints *n* = 1281) (left panels). ROC curves depict prognostic performance for incipient TB, stratified by time intervals to disease (< 3, < 6, <12, <18, < 24, < 30 months) (middle panel) and mutually exclusive time intervals to disease (0-3, 3-6, 6-12, 12-18, 18-24, 24-30 months) (right panel). AUC and 95% confidence intervals for each interval to disease are shown.

**Supplementary Figure 10**. *Continued*

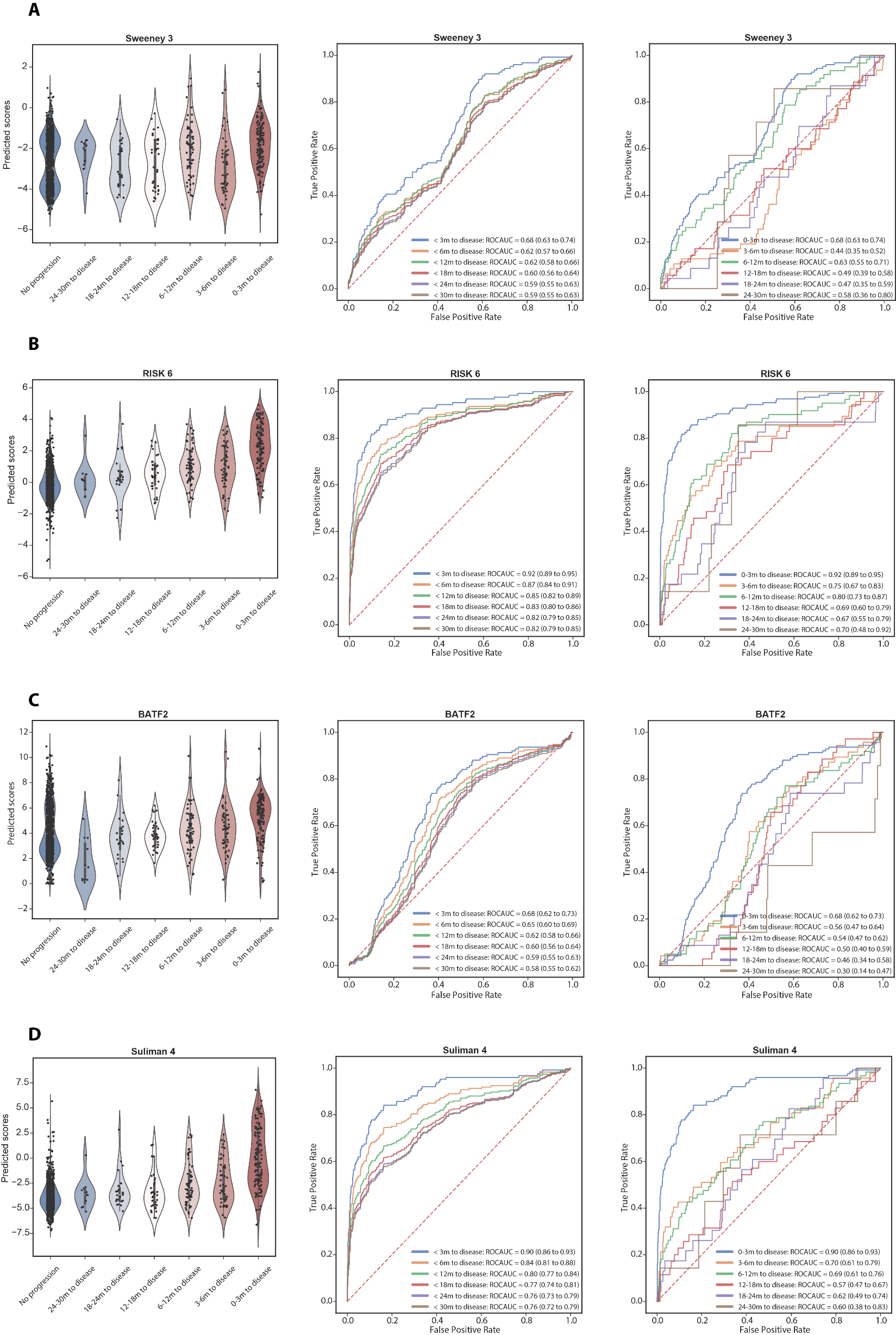

**Supplementary Figure 10**. *Continued*

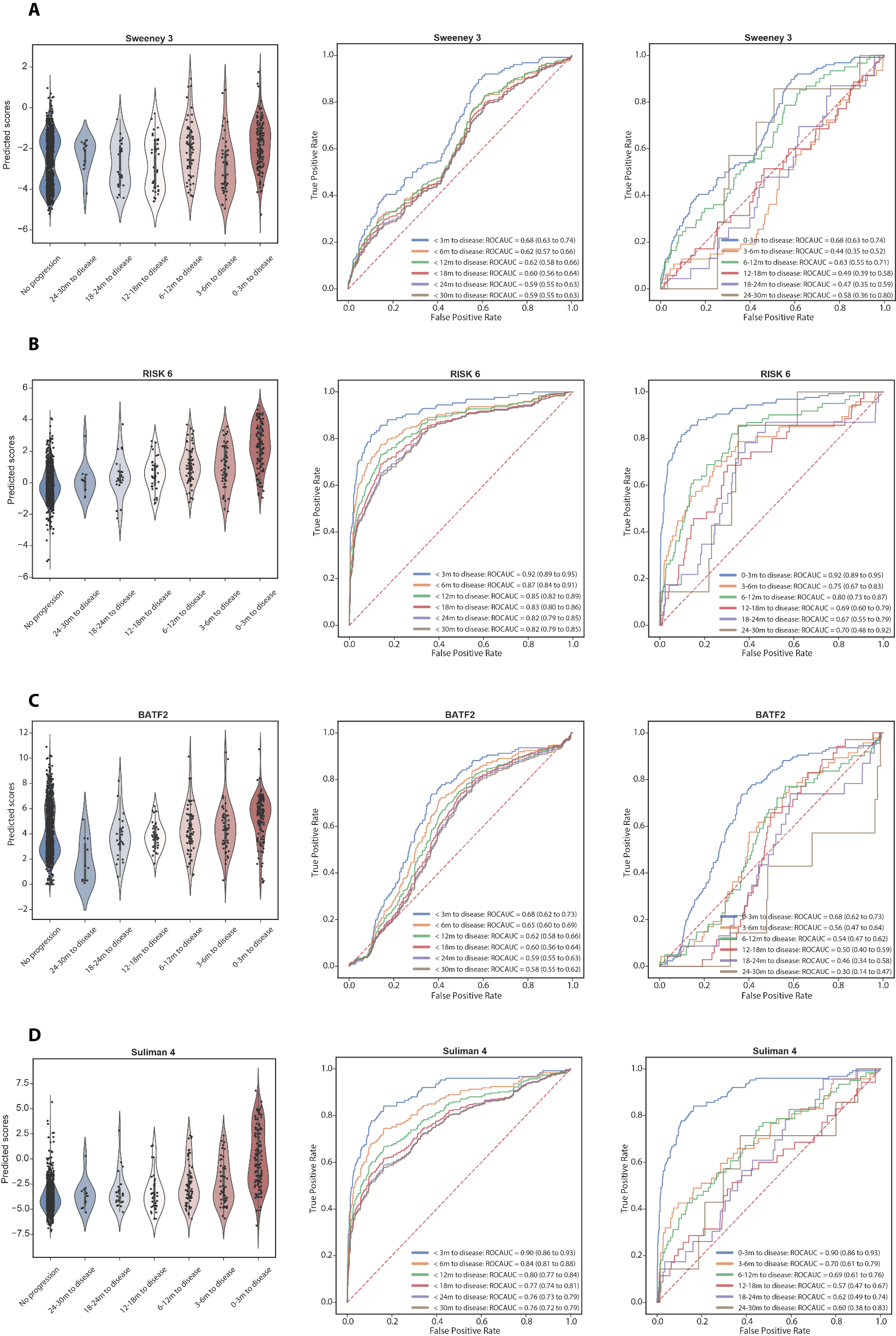

#### Supplementary Figure 11

Diagnostic performance of our models and the 4 published models(*3-6*) for active TB using a pooled dataset of all 57 collected cohort studies (**Supplementary Table. 1**) (datapoints *n* = 6290). The distributions of TB scores, stratified by different TB disease stages (HC, LTBI, and ATB), other lung disease (OLD) and viral infection (VI), are visualized in the violin plots (A, C, E, G, I, and K). ROC curves (B, D, F, H, J and L) depict diagnostic performance of the models. AUCs and 95% confidence intervals for each comparison are also shown.

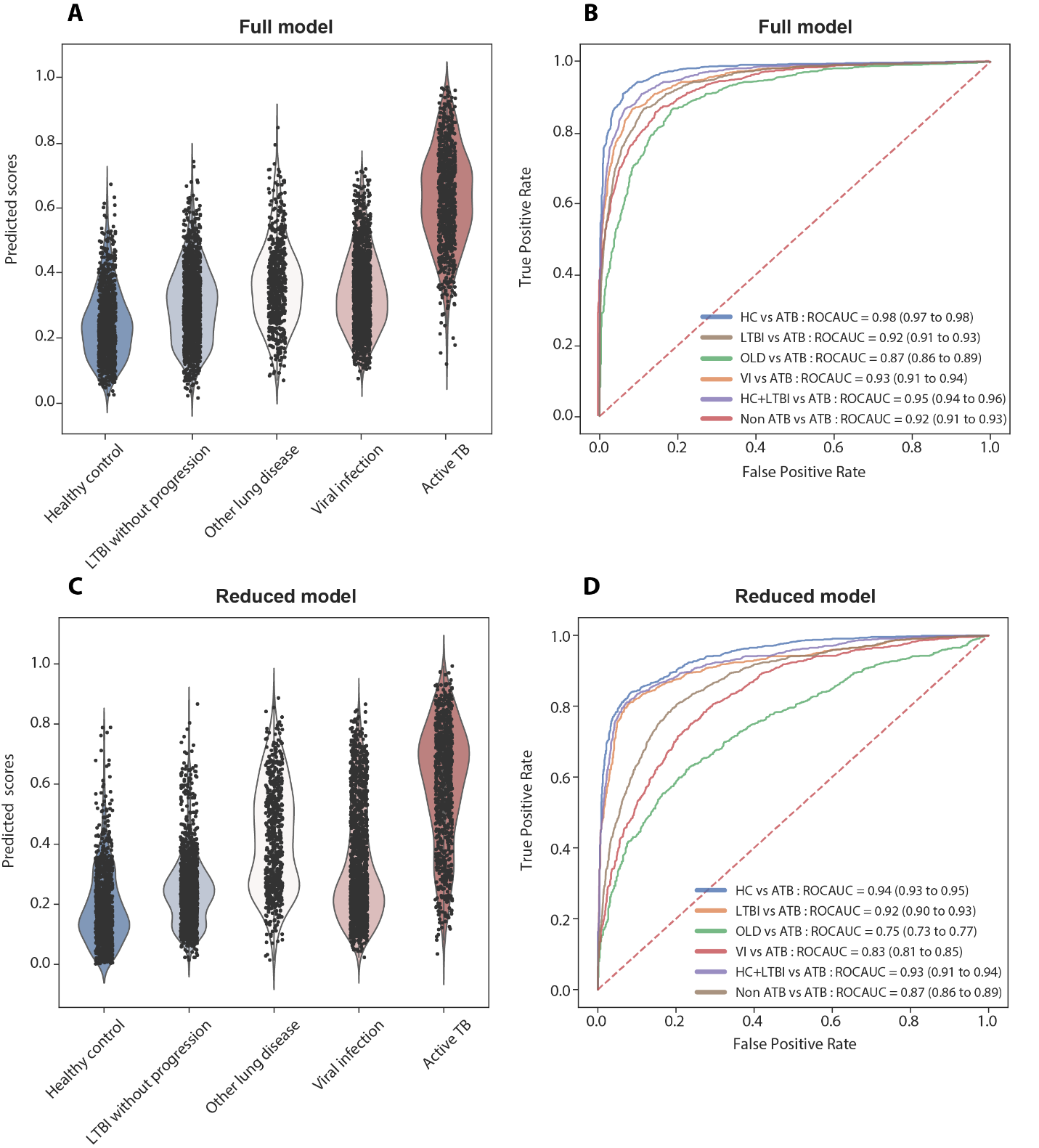

**Supplementary Figure 11**. *Continued*

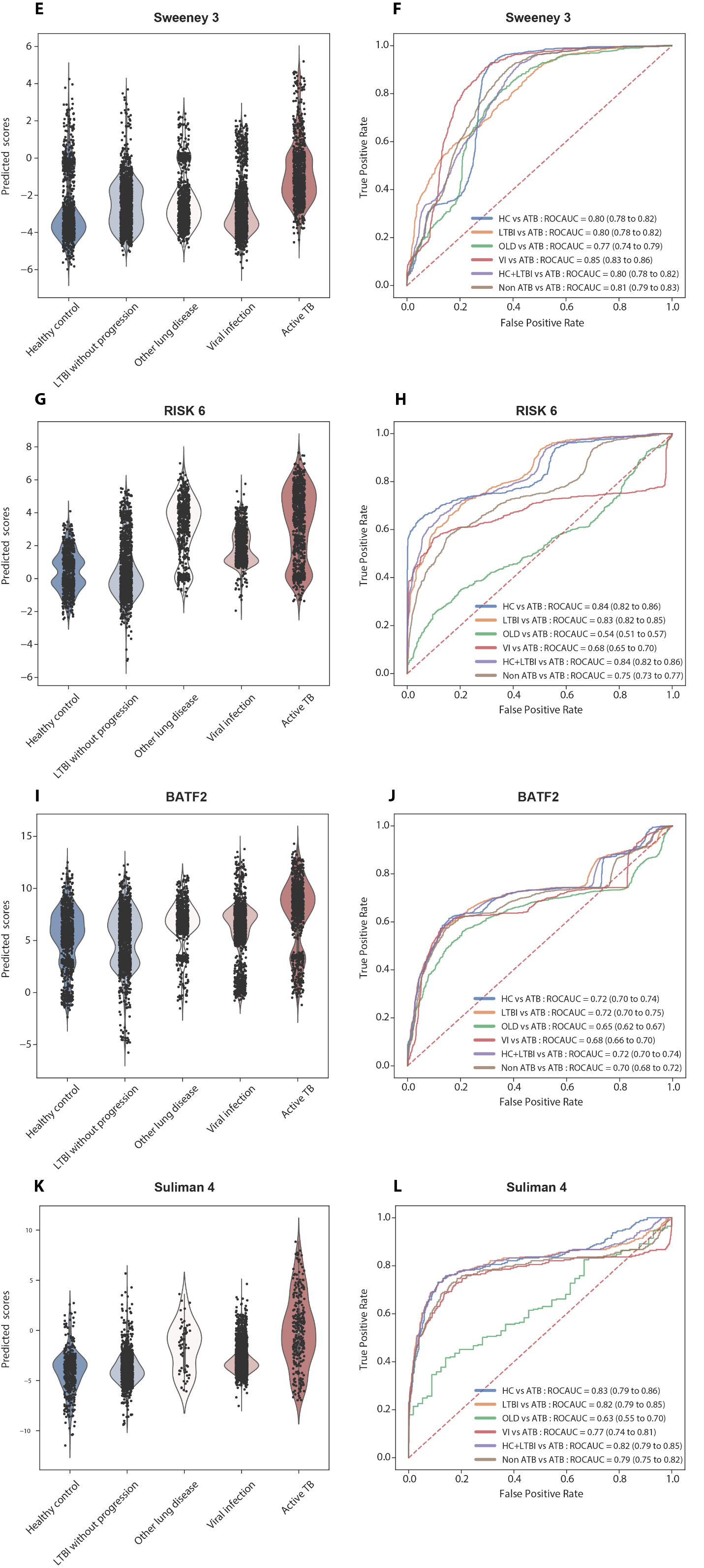

**Supplementary Figure 11**. *Continued*

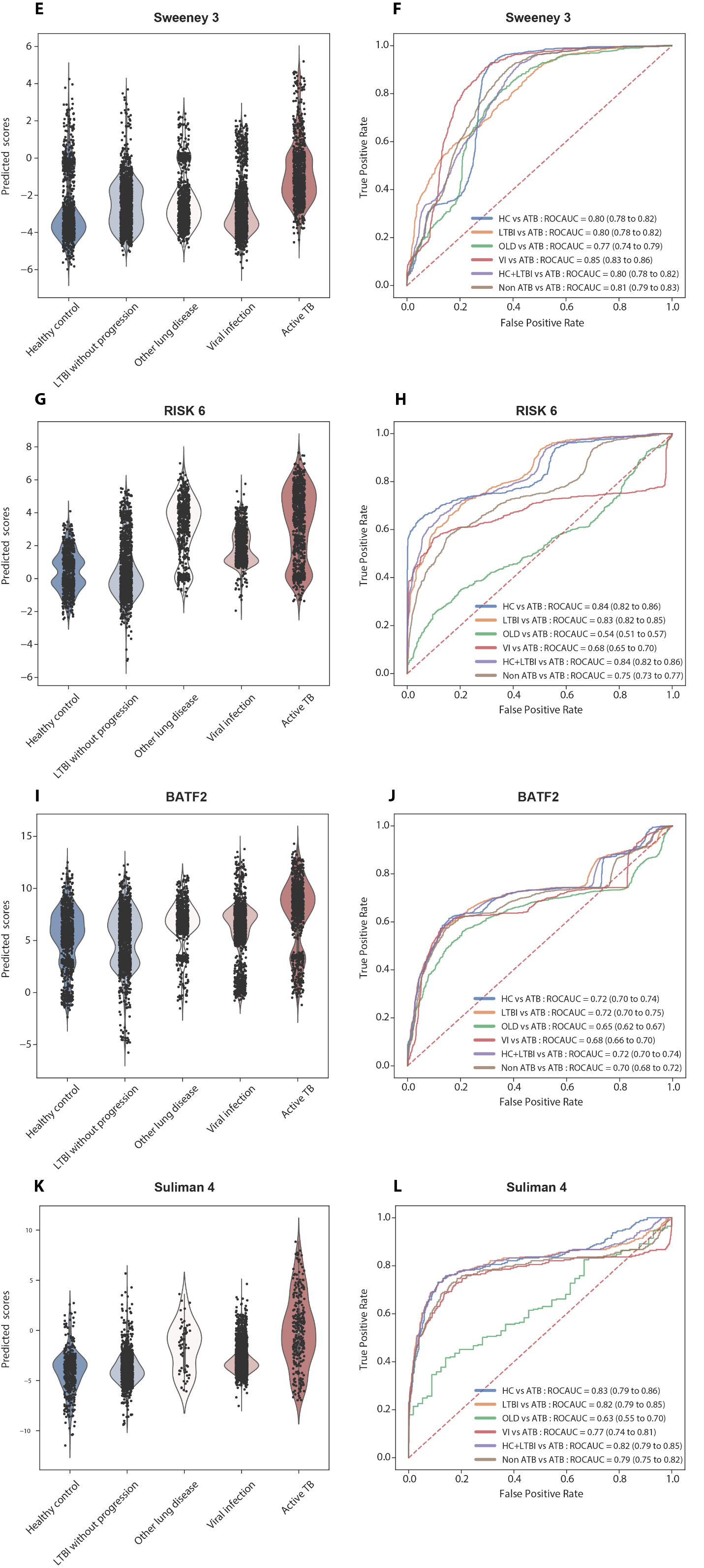

#### Supplementary Figure 12

The temporal dynamics of TB scores generated by the reduced model, stratified by the time of sputum culture conversion to negative (negativity at day 28, 56, 84 and 168, and no conversion at day 168 [failed]). The dashed line represents the TB scores of each patient responding to treatment over time, and the red line represents the median of the stratified group with 95% confidence interval shown in the shaded area.

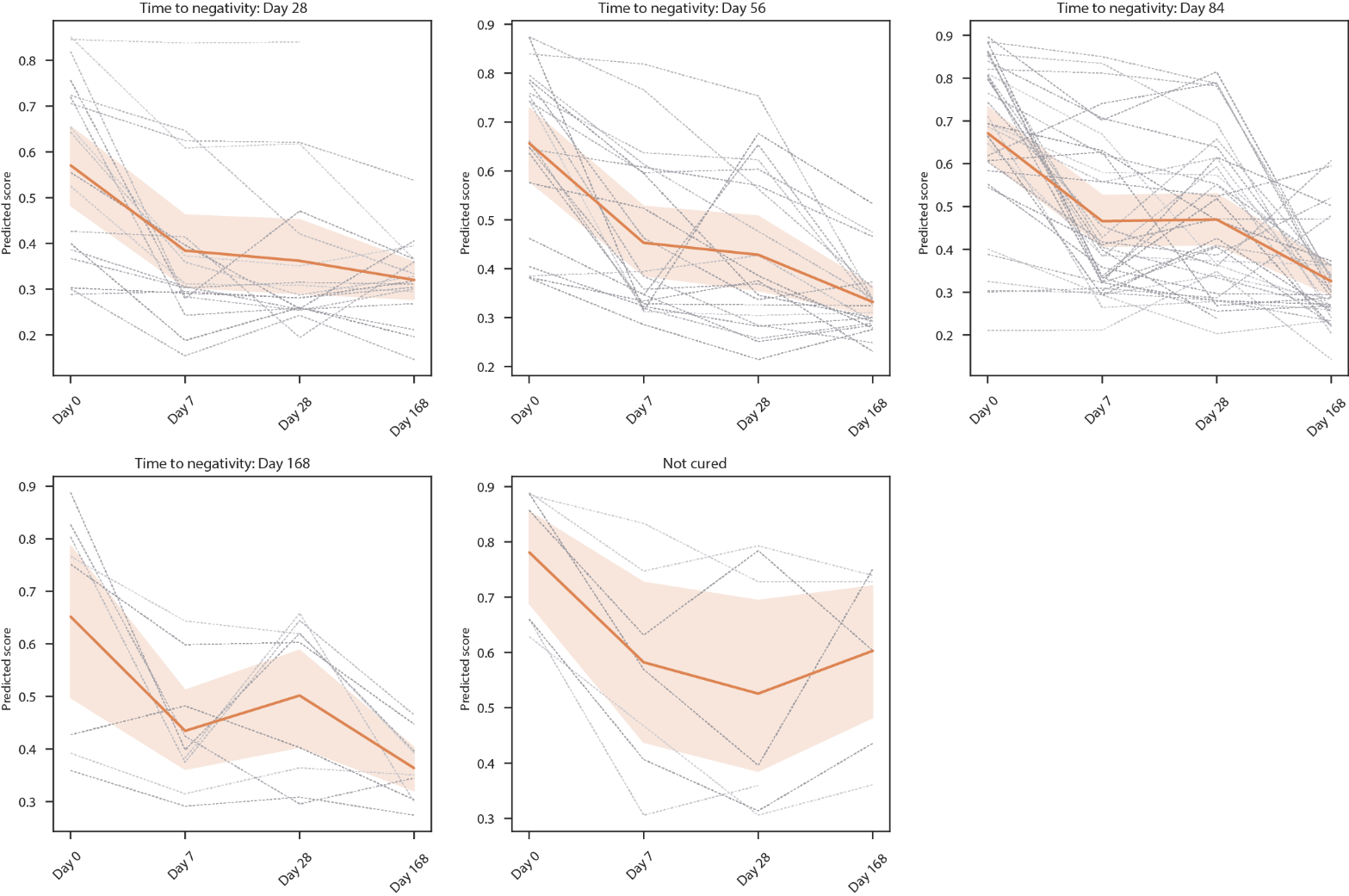

#### Supplementary Figure 13

Predictive performance of the reduced model in recurrent TB disease. ROC curves depicting predictive performance of the model for discrimination between cured patients with or without TB recurrence within 2 years after treatment completion are shown in different colors, stratified by different timepoints after treatment initiation. AUC and 95% confidence intervals for each interval to disease are also shown.

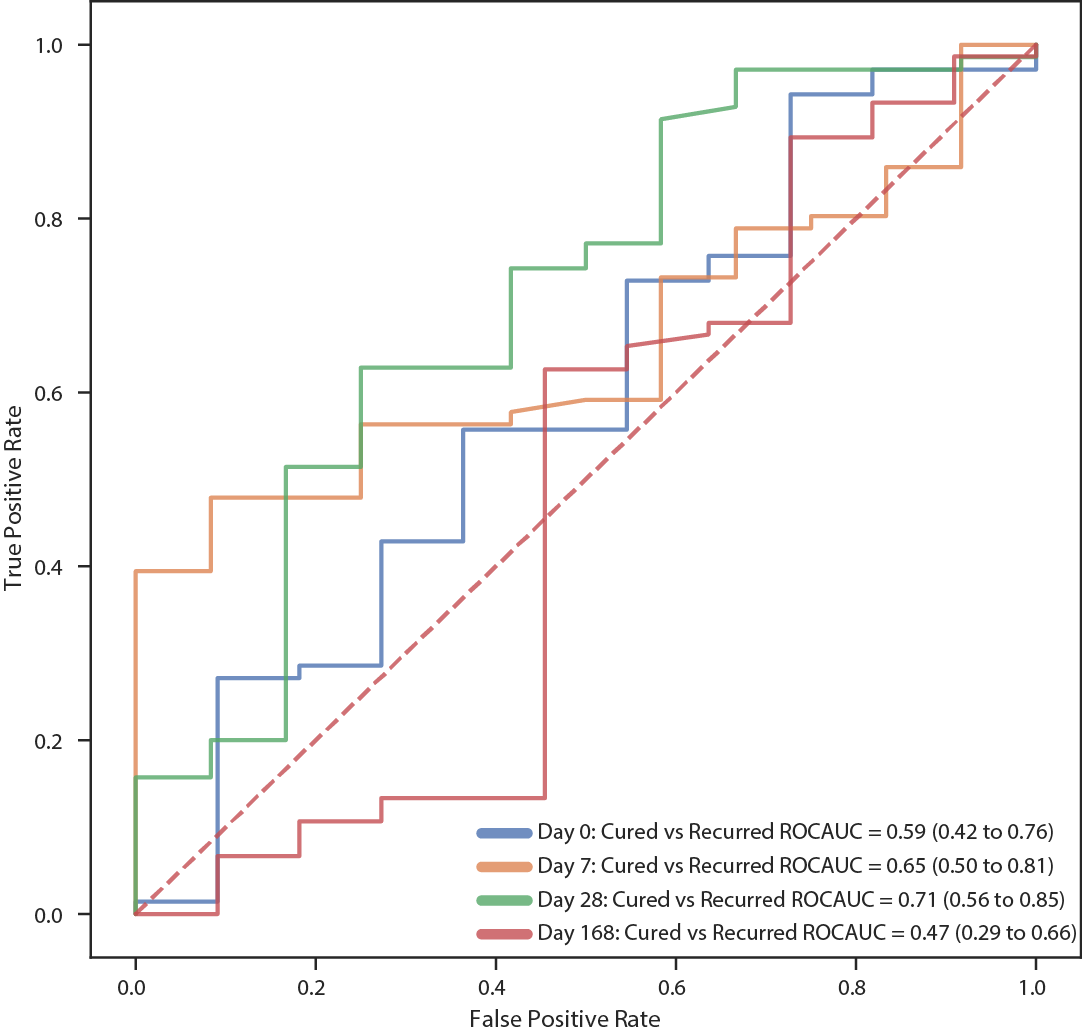

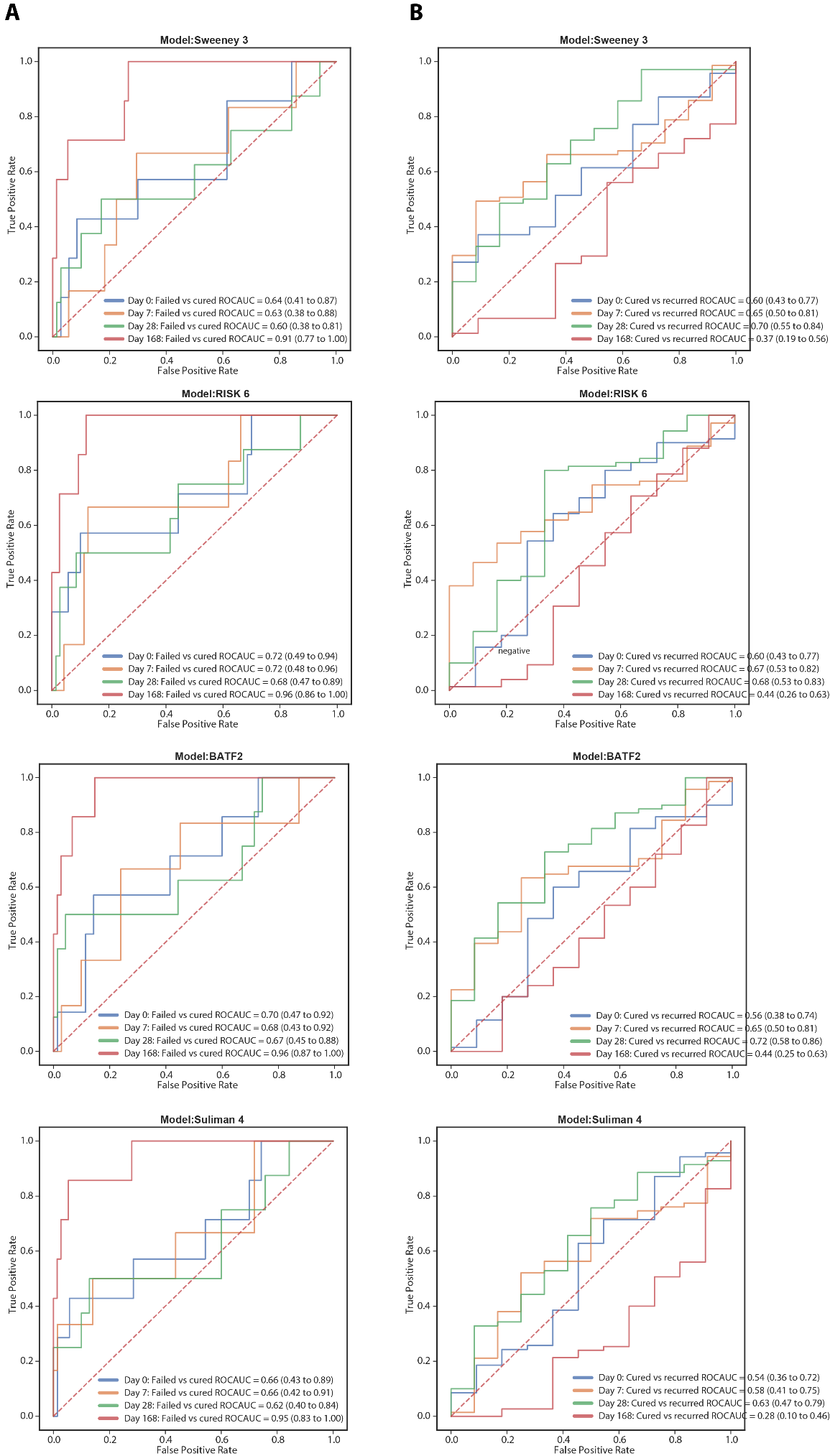

#### Supplementary Figure 14

Predictive performance of the 4 published models. ROC curves, stratified by different timepoints after treatment initiation, depict predictive performance of the models for discrimination between patients with bacteriological cure and those with treatment failure at EOT (A) and cured patient with or without TB recurrence within 2 years after treatment completion (B). AUC and 95% confidence intervals for each interval to disease are also shown.

#### Supplementary Figure 15

The relationship between TB scores generated by the reduced model and *in vivo* pulmonary inflammation. The Spearman correlations between TB scores and MGIT culture time to positivity (A), Xpert Ct values (B), total glycolytic ratio activity (TGRA) at three time points after treatment initiation (Day 0, Day 28 and Day168) (C-E) are displayed. The scores at baseline stratified by radiologically persistent or cleared lung inflammation at EOT are shown in the violin plot (F).

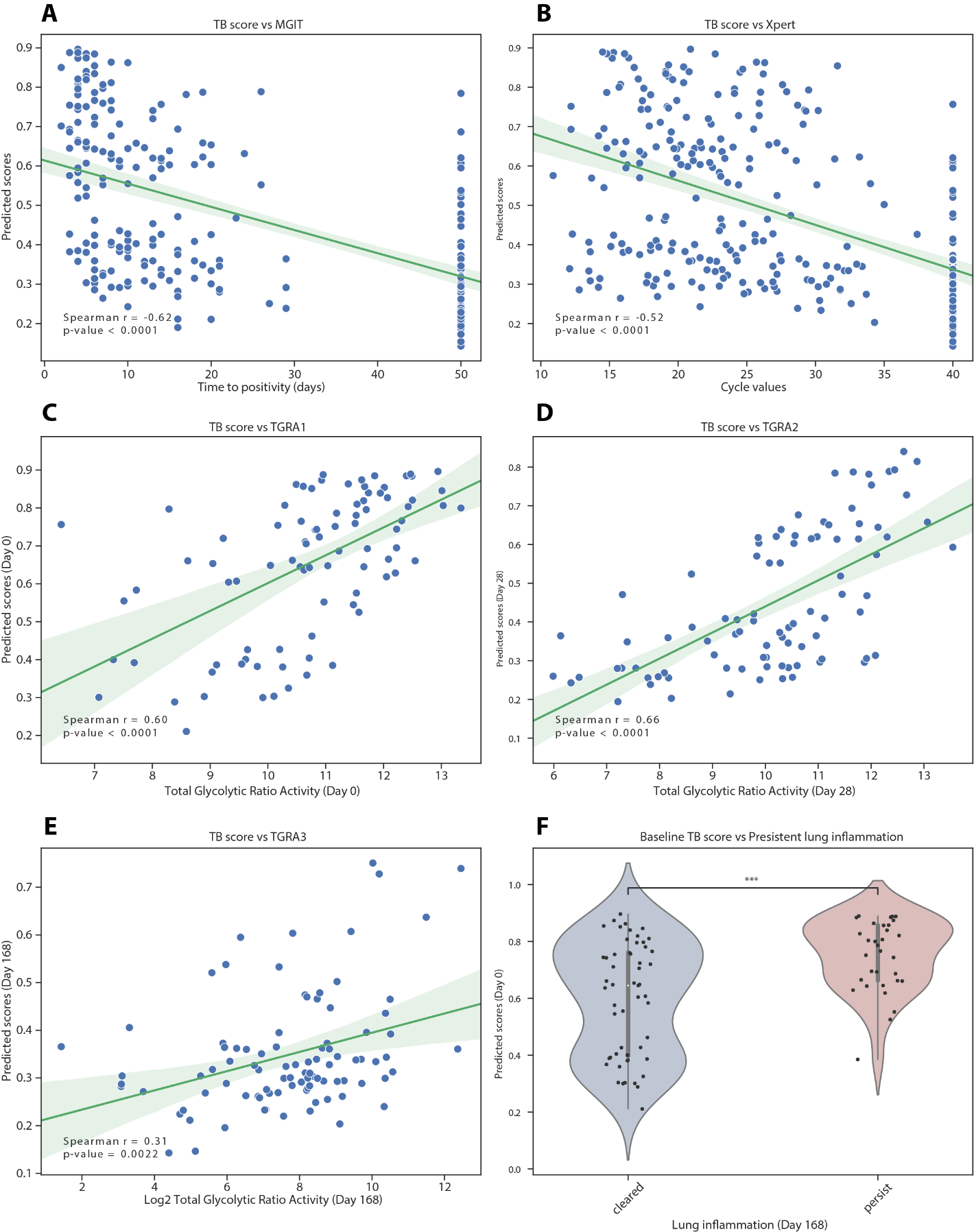

### REFERENCES

1.	R. K. Gupta *et al.*, Concise whole blood transcriptional signatures for incipient tuberculosis: a systematic review and patient-level pooled meta-analysis. *The Lancet Respiratory Medicine* **8**, 395-406 (2020).

2.	H. Warsinske, R. Vashisht, P. Khatri, Host-response-based gene signatures for tuberculosis diagnosis: A systematic comparison of 16 signatures. *PLoS Med* **16**, e1002786 (2019).

3.	T. E. Sweeney, L. Braviak, C. M. Tato, P. Khatri, Genome-wide expression for diagnosis of pulmonary tuberculosis: a multicohort analysis. *Lancet Respir Med* **4**, 213-224 (2016).

4.	A. Penn-Nicholson *et al.*, RISK6, a 6-gene transcriptomic signature of TB disease risk, diagnosis and treatment response. *Sci Rep* **10**, 8629 (2020).

5.	J. Roe *et al.*, Blood Transcriptomic Stratification of Short-term Risk in Contacts of Tuberculosis. *Clin Infect Dis* **70**, 731-737 (2020).

6.	S. Suliman *et al.*, Four-gene Pan-African Blood Signature Predicts Progression to Tuberculosis. *Am J Respir Crit Care Med*, (2018).
